# Interactions with multiple inner kinetochore proteins determine mitotic localization of FACT

**DOI:** 10.1101/2024.06.14.599021

**Authors:** Julia Schweighofer, Bhagyashree Mulay, Ingrid Hoffmann, Doro Vogt, Marion E. Pesenti, Andrea Musacchio

## Abstract

The FAcilitates Chromatin Transcription (FACT) complex is a dimeric histone chaperone that operates on chromatin during transcription and replication. FACT also interacts with a specialized centromeric nucleosome containing the histone H3 variant CENP-A and with CENP-TW, two subunits of CCAN, a 16-protein complex associated with CENP-A. The significance of these interactions remains elusive. Here, we show that FACT has multiple additional binding sites on CCAN. The interaction with CCAN is strongly stimulated by casein kinase II (CK2) phosphorylation of FACT. Mitotic localization of FACT to kinetochores is strictly dependent on specific CCAN subcomplexes. Unexpectedly, we also find that DNA readily displaces FACT from CCAN, suggesting that FACT becomes recruited through a pool of CCAN that is not stably integrated into chromatin. Collectively, our results point to a potential role of FACT in chaperoning CCAN during transcription or in the stabilization of CCAN at the centromere during the cell cycle.

**Teaser:** DNA-sensitive, direct interactions with multiple inner kinetochore subunits deliver FACT to the kinetochore.

## Introduction

Chromosomes are DNA packaging structures that consist of a single molecule of DNA and many different associated proteins. They contain several functionally specialized regions that work in conjunction with transcription, replication, and inheritance. A notable specialized chromatin locus is the centromere. The histone H3 variant centromere protein A (CENP-A) is greatly enriched at centromeres and is considered the crucial epigenetic marker of centromeres. CENP-A seeds the kinetochore, a large protein complex that connects the replicated chromosomes (sister chromatids) to spindle microtubules during mitosis to ensure their equal distribution to the daughter cells (*1*, *2*). Its presence at centromeres recruits specialized machinery that delivers new CENP-A at every cell cycle to compensate for its dilution during DNA replication (*3*).

The kinetochore is divided into inner and outer layers (*4*). The outer layer, consisting of 10 proteins collectively referred to as the Knl1 complex, Mis12 complex, Ndc80 complex (KMN) network and associated proteins is assembled during mitosis to directly attach to spindle microtubules (*5*). The inner layer, consisting of 16 proteins collectively referred to as the constitutive centromere associated network (CCAN), bridges the centromeric chromatin and outer kinetochore and localizes to the centromere throughout the cell cycle (*6*, *7*). The CCAN consists of different subunits and subcomplexes, including CENP-C, CENP-HIKM, CENP-LN, CENP-OPQUR, and CENP-TWSX (*8*) (Fig. 1A).

**Figure 1.**
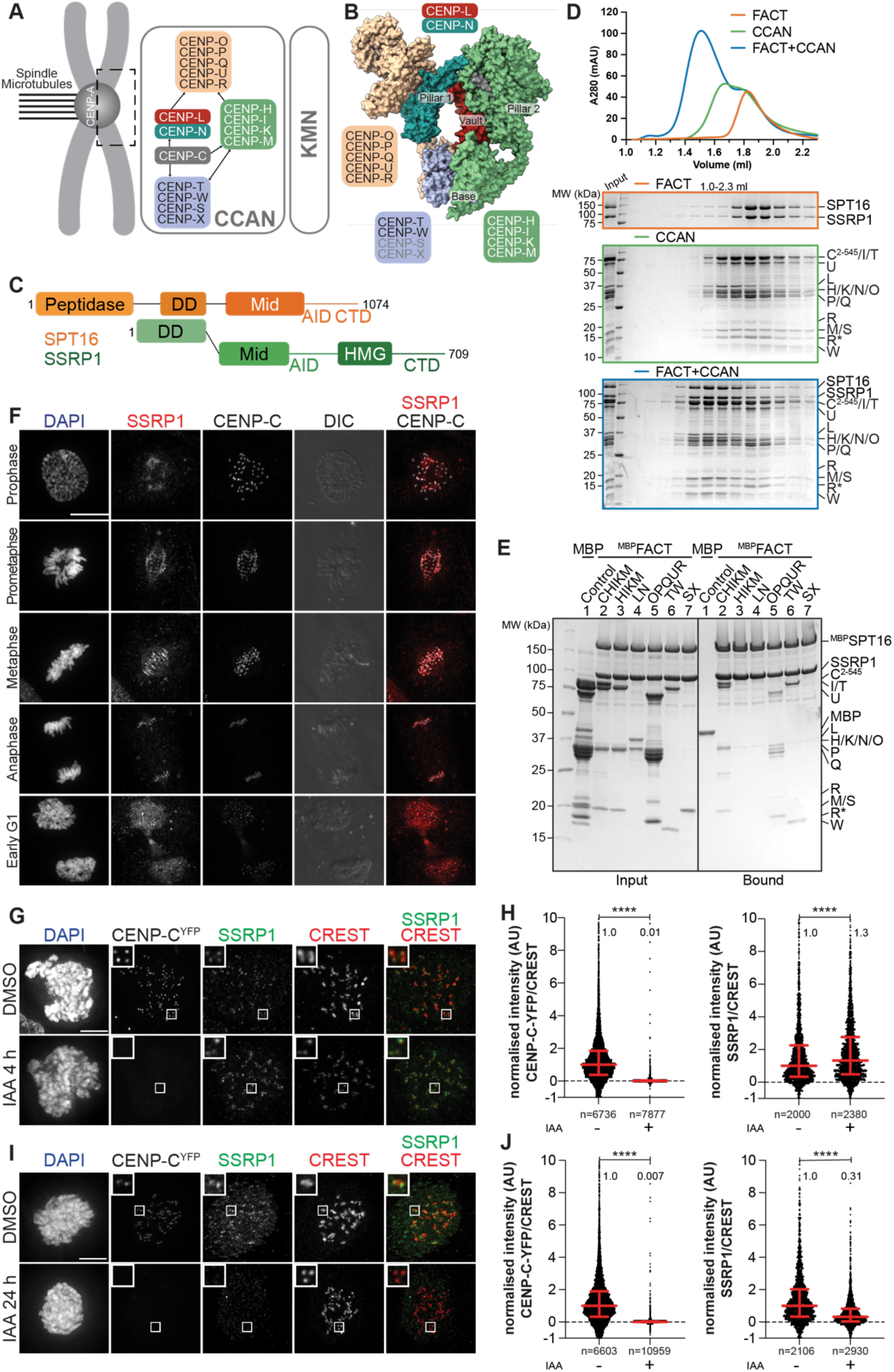
FACT forms a complex with the CCAN *in vitro* and mitotic localization at the kinetochore depends on the CCAN. (**A**) Scheme of the human kinetochore with focus on subunits of the CCAN. (**B**) Surface representation of the structure of the human CCAN based on PDB 7QOO with the subcomplexes colored as in (A). The structure lacks subunits CENP-SX. (**C**) Scheme of the domain architecture of the FACT complex, SPT16 in orange, SSRP1 in green. Peptidase, peptidase-like domain; DD, dimerization domain; Mid, Mid domain; AID, acidic intrinsically disordered domain; CTD, C-terminal domain; HMG, high mobility group domain. (**D**) Analytical size exclusion chromatography (SEC) of FACT, CCAN including CENP-C^2-545^, and the resulting 18-subunit complex. Fractions were analyzed by SDS-PAGE and visualized by Coomassie staining. (**E**) Amylose-resin pull-down assay with FACT, where SPT16 has an N-terminal MBP-tag, immobilized on beads and isolated CCAN subcomplexes as preys. (**F**) Representative images of localization of SSRP1 during mitosis. Asynchronous RPE-1 cells were immunostained for SSRP1 to visualize FACT complex, CENP-C to visualize the kinetochores, and DAPI to stain DNA. Scale bar: 5 μm. (**G**) Representative images of localization of SSRP1 4 hours after addition of IAA to degrade CENP-C in DLD-1-CENP-C^YFP-AID^ cells. Cells were treated with IAA (500µM) to degrade endogenous CENP-C and Nocodazole (3.3 µM) for 4 hours to enrich for mitotic cells. CREST serum was used to visualize kinetochores and DAPI to stain DNA. Three biological replicates were performed. Scale bar: 5 μm. (**H**) Scatter plots of YFP and SSRP1 levels at kinetochores for the experiment shown in panel (G). *n* is the number of individual kinetochores measured. (**I**) Representative images of localization of SSRP1 24 hours after addition of IAA to degrade CENP-C in DLD-1-CENP-C^YFP-AID^ cells. Cells were treated with IAA (500 µM) to degrade endogenous CENP-C for 24 hours and Nocodazole (3.3 µM) for 4 hours to enrich for mitotic cells. CREST serum was used to visualize kinetochores and DAPI to stain DNA. Three biological replicates were performed. Scale bar: 5 μm. (**J**) Scatter plots of YFP and SSRP1 levels at kinetochores for the experiment shown in panel (I). *n* is the number of individual kinetochores measured.

Two CCAN proteins, CENP-C and CENP-N, decode the centromere by recognizing CENP-A (*9–15*). In addition to binding CENP-A, CENP-C interacts directly with other inner kinetochore subunits, including CENP-HIKM and CENP-LN as well as the outer kinetochore complex MIS12 (*12*, *16–20*). A second subunit, CENP-T, binds stably to CENP-W and connects the CCAN and the outer kinetochore by interacting with Mis12 and Ndc80 complexes (Mis12C and Ndc80C, respectively) through its long disordered N-terminal tail (*21*, *22*). CENP-W and the C-terminal region of CENP-T consist of a histone fold domain (HFD). The CENP-TW subcomplex further tetramerizes with two additional HFD-containing proteins, CENP-S and CENP-X. It has been reported that the resulting CENP-TWSX complex is integrated into centromeric chromatin as a nucleosome-like particle (*23*, *24*). Recent structural work has shown that CCAN consists of two structural pillars (composed of CENP-HIKM and CENP-OPQUR) flanking a central DNA-binding vault (contributed by CENP-LN) and a base (CENP-TWSX; Fig. 1B). The central vault enables tight binding of linker DNA by CCAN. *In vitro* and *in vivo*, CENP-A has been shown to form an octameric nucleosome consisting of a CENP-A/H4 tetramer flanked by two H2A/H2B dimers wrapped by ∼150 base pairs (bp) DNA (*25*). The CENP-A nucleosome has been proposed to neighbour the CCAN structure bound to linker DNA (*26–28*).

The original CENP-A co-precipitation experiments that identified CCAN subunits also identified the FAcilitates Chromatin Transcription (FACT) complex for a specific interaction with CENP-A nucleosomes (*6*, *29–32*). FACT is a H2A/H2B chaperone that prevents histone loss whilst facilitating assembly and disassembly of nucleosomes during transcription (*33–36*). Additionally, it has been implicated in DNA replication and repair (*37–46*). FACT is a heterodimer of Suppressor of Ty protein 16 (SPT16) and Structure-specific recognition protein 1 (SSRP1), both large multi-domain proteins with an array of Plekstrin homology (PH) domains (*47–49*). SPT16 has an N-terminal peptidase-like domain, which has lost its catalytic activity but interacts with Mini Chromosome Maintenance protein complex (MCM) 2-7 and with the fork protection complex during replication, as well as with the Set3 histone deacetylase complex (*50–52*). SPT16 Mid domain binds to histone H3/H4 tetramers. The subsequent acidic intrinsically disordered (AID) segment associates with H2A/H2B dimers (*53–55*). SSRP1 contains an high mobility group (HMG) domain, which is associated with DNA binding (*56*, *57*) (Fig. 1C).

The precise significance of the interaction of FACT with centromeres remains elusive. In chicken, *Drosophila*, budding and fission yeast, FACT has been implicated in CENP-A deposition and in preventing ectopic localization of CENP-A (*58–61*). In humans, FACT has been shown to directly interact with CENP-TW HFDs via the AID of SPT16 (*62*). In this study, we demonstrate that the interaction of FACT with CCAN is complex, with additional binding sites on CENP-C and CENP-OPQUR. FACT engages in a stable 18-subunit complex with CCAN, whose assembly requires the phosphorylation of FACT by the constitutively active kinase casein kinase II (CK2). Mitotic localization of FACT at the kinetochore is dominated by CENP-HIKM and CENP-TW. We find that DNA displaces FACT from CCAN, suggesting a potential role of FACT in chaperoning CCAN during transcription or in the deposition of CCAN at the centromere during or after replication.

## Results

### FACT forms a stable complex with CCAN *in vitro*

As CCAN and FACT co-precipitate with CENP-A nucleosomes and FACT has been proposed to bind directly to CCAN subunits, we asked if a CCAN/FACT complex could be reconstituted *in vitro* using recombinant proteins. Previously, we have reconstituted a 16-subunit CCAN from four stable subcomplexes, including CENP-CHIKM (assembled with C^2-545^, i.e. with a fragment of CENP-C encompassing residues 2-545), CENP-LN, CENP-OPQUR and CENP-TWSX (*26*, *63*, *64*). We reconstituted CCAN starting from these subcomplexes (Fig. S1A). In analytical size-exclusion chromatography (SEC) experiments, FACT and CCAN co-eluted in a single peak and at earlier elution volumes relative to the individual complexes, indicating that CCAN and FACT bind directly in an 18-subunit complex (Fig. 1D).

To identify CCAN subunits involved in FACT binding, we immobilized FACT on amylose-resin through an N-terminal MBP-tag on SPT16 (^MBP^FACT), and used the various CCAN subcomplexes as preys. In addition to confirming the previously reported interaction with CENP-TW (*62*), we observed interactions with CENP-OPQUR and CENP-C^2-545^HIKM (Fig. 1E). The latter interaction required CENP-C^2-545^, because the CENP-HIKM complex, which lacks CENP-C, did not bind (Fig. 1E, lane 3), as opposed to previously published experiments with avian CCAN, which suggest a direct interaction with CENP-HIKM (*58*). CENP-SX, which contains HFDs similar to CENP-TW (*24*), did not bind to ^MBP^FACT (Fig. 1E, lane 7). We confirmed the association of FACT with CENP-TW and CENP-OPQUR by analytical SEC, whereas FACT and CENP-C^2-545^HIKM did not form a stable complex in solution (Fig. S1B). We conclude that FACT and CCAN bind directly, and that the interaction is mediated by multiple binding interfaces.

### Mitotic localization of FACT to the kinetochore depends on the CCAN

FACT localizes to chromatin, especially nucleoli, in interphase, reflecting its role in transcription (*65*, *66*). In chicken DT40 cells, FACT was also observed to localize at centromeres during mitosis (*58*). To investigate mitotic FACT localization in human cells, we stained SSRP1 by immunofluorescence in hTERT-immortalized retinal pigment epithelial (RPE-1) cells. FACT localized to the kinetochore in all mitotic phases, exhibiting a more diffuse signal in early and late mitosis (Fig. 1F). To dissect how FACT is recruited to kinetochores during mitosis, we exploited a previously described colorectal adeno-carcinoma DLD-1 cell for rapid degradation of CENP-C (*67*). In this system, both CENP-C alleles are endogenously tagged with an auxin-inducible degron (AID) (*68*) and an enhanced yellow fluorescent protein (EYFP). After a 4-hour treatment of cells arrested in mitosis by nocodazole treatment with the auxin derivative indole acetic acid (IAA), CENP-C was completely depleted from kinetochores. Instead, CENP-HK and CENP-TW localization was largely unaffected, as previously observed (*26*). SSRP1 localization was also largely unaffected (Fig. 1G,H and Fig. S2), indicating that recruitment of FACT is independent of CENP-C or that FACT remains stably localized after initial depletion of CENP-C. When the treatment with IAA was extended to 24 hours, however, the kinetochore levels of FACT were greatly decreased. This correlated with modest to strong decreases in CCAN subunit localization (Fig. 1I,J and Fig. S3). Of note, these experiments also revealed that the course of depletion of CCAN subunits after CENP-C degradation in mitotically arrested cells does not completely recapitulate the pattern of depletion seen in cycling cells (*26*) (Fig. S2 and Fig. S3). Collectively, these observations link kinetochore localization of FACT to the interactions with CCAN observed *in vitro*, although they do not exclude a potential role of centromere transcriptional activity in the recruitment and retention of FACT during mitosis (*69–72*).

### Cooperative and anti-cooperative FACT/CCAN binding

To further characterize how individual interactions of CCAN and FACT stabilize their assembly, we titrated CCAN subcomplexes in different combinations in an *in vitro* pull-down assay with ^MBP^FACT as bait (Fig. 2A). We quantified the results using the band intensities of CENP-M, CENP-U, CENP-L and CENP-W, which were well resolved in SDS-PAGEs, as representative of their cognate CCAN subcomplexes (Fig. 2B-E). As shown above (Fig. 1E), CENP-C^2-545^HIKM and CENP-OPQUR bound FACT (Fig. 2A). A CENP-TW complex consisting only of the histone fold domain (HFD) of these proteins (CENP-T^458-C^/full-length CENP-W, henceforth CENP-TW^HFD^) also bound FACT (Fig. 2A, lane 4). However, CENP-M, CENP-U and CENP-W exhibited a markedly lower band intensity when their cognate subcomplex was exposed to FACT without the other CCAN subcomplexes (Fig. 2A-E, lanes 2-4). When exposed to additional subcomplexes (lanes 5-12), stronger binding was observed. Notably, addition of CENP-C^2-545^ to HIKM (instead of isolated CENP-HIKM) enhanced binding when certain subcomplexes were omitted (e.g. CENP-TW in lanes 5, 6; CENP-OPQUR and CENP-LN in lanes 7, 8; and CENP-OPQUR in lanes 9, 10). However, when the complete CCAN was used as prey, absence of CENP-C^2-545^ did not significantly change the level of bound CCAN subunits (lanes 11,12). Collectively, these results are consistent with the idea that FACT binding involves multiple interaction interfaces of CCAN.

**Figure 2.**
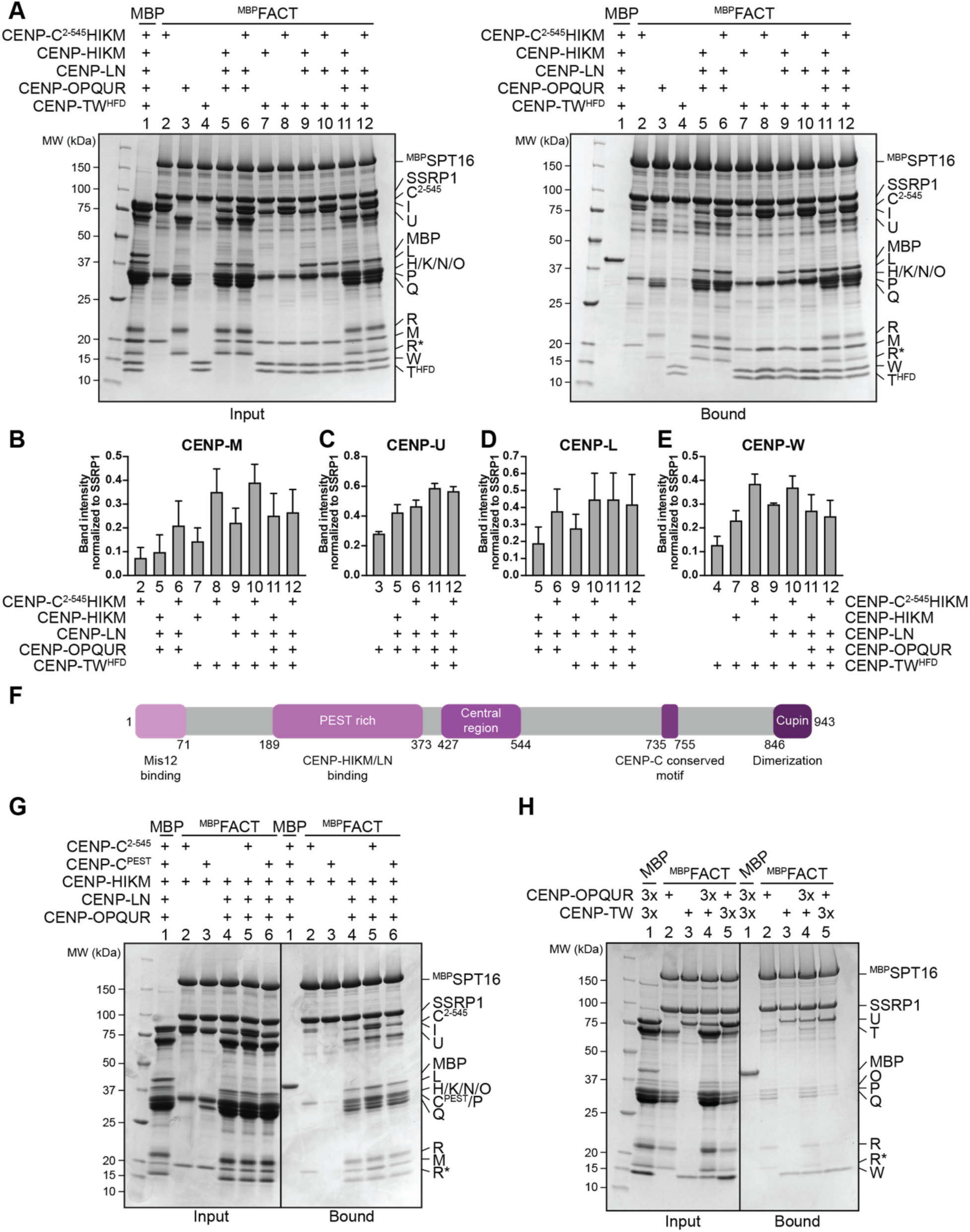
CCAN binds FACT cooperatively. **(A)** Amylose-resin pull-down assay with ^MBP^FACT as bait and adding different combinations of CCAN subcomplexes as preys as indicated above the SDS-PAGE gels. MBP was used as a negative control. (B-E) Quantifications of the pull-down in panel (A) from three repeats. The band intensity of the target protein was normalized to SSRP1. One subunit per subcomplex was quantified: (**B**) CENP-M, (**C**) CENP-U, (**D**) CENP-L, (**E**) CENP-W. (**F**) Scheme of CENP-C with functional domains and their residue number indicated. (**G**) Amylose-resin pull-down assay with ^MBP^FACT as a bait to assess the binding of CENP-HIKM or CENP-HIKMLNOPQUR in presence of CENP-C^2-545^ or CENP-C^PEST^. (**H**) Amylose-resin pull-down assay with ^MBP^FACT as a bait and either CENP-OPQUR or -TW added in molar excess.

CENP-C binds to CENP-HIKMLN through its PEST (proline-glutamic acid-serine-threonine)-rich region (CENP-C^189-400^, CENP-C^PEST^) (*12*, *16*, *18*, *19*) (Fig. 2F). CENP-C^PEST^, however, was neither capable of a direct interaction with FACT when combined with CENP-HIKM, nor did it trigger increased binding of CENP-HIKMLNOPQUR to ^MBP^FACT (Fig. 2G). These observations suggested that CENP-C and FACT may bind directly outside of the CENP-C^PEST^. To identify regions of CENP-C involved in FACT binding, we divided the sequence of CENP-C in different fragments, expressed them as fusions to MBP, and used them as baits in a pull-down assay. FACT bound CENP-C^401-545^, CENP-C^401-600^, CENP-C^546-600^, CENP-C^721-759^, and CENP-C^721-943^ (Fig. S4A), which collectively encompass 1) the CENP-C central region, 2) a region adjacent to the central region, 3) the CENP-C conserved motif, and 4) the C-terminal cupin domain involved in dimerization. As both the central region and the CENP-C conserved motif bind specifically to CENP-A nucleosomes (CENP-A nucleosome core particle, CENP-A^NCP^), these observation suggest that FACT may stabilize the CENP-A nucleosome binding region of CENP-C in the absence of nucleosomes (Fig. 2F) (*9*, *11*, *14*, *15*). Confirming this conclusion, inclusion of CENP-A^NCP^ in a pull-down assay where FACT was bound to immobilized ^MBP^CENP-C^EGFP^ (a full-length CENP-C construct) caused FACT to dissociate (Fig. S4B), indicating that binding of FACT and nucleosomes to CENP-C is mutually exclusive.

An unexpected aspect of the CCAN interaction with FACT is that the addition of CENP-OPQUR appeared to reduce the levels of CENP-HIKM and CENP-TW (using CENP-M and CENP-W as readouts, respectively; Fig. 2A lanes 11, 12 and quantified in Fig. 2B and E). Within CCAN, CENP-OPQUR and CENP-TW do not directly bind to each other and require CENP-LN and CENP-HIKM for their interaction (*26*, *27*, *63*). As they are both able to bind FACT, however, we anticipated that FACT may bridge these complexes. Contrary to this expectation, CENP-OPQUR and CENP-TW competed for FACT, with CENP-TW showing a higher affinity for FACT (Fig. 2H).

To investigate this phenomenon further, we tried to shed light on the determinants of the interaction of FACT with CENP-OPQUR. We found the disordered N-terminal tails of CENP-Q and CENP-U to be required for CENP-OPQUR/FACT binding, because a truncation of these tails (CENP-OPQ^68-C^U^115-C^R, herewith indicated as CENP-OPQ^ΔN^U^ΔN^R) completely abolished the association with ^MBP^FACT (Fig. S4C). This result was confirmed *in vivo*, where FACT was identified in immunoprecipitates (IPs) of ^EGFP^CENP-U but not of CENP-U^115-C^ (Fig. S4D). However, CENP-C^2-545^HIKMLNOPQ^ΔN^U^ΔN^R bound ^MBP^FACT, probably because CENP-C^2-545^ provides sufficient binding affinity for the FACT complex. Even though CENP-OPQ^ΔN^U^ΔN^R does not bind FACT, it continued to oppose binding of FACT to CENP-TW (Fig. S4C), possibly through an allosteric mechanism.

### FACT localization to the kinetochore requires CENP-HIKM and-TW

As the mitotic localization of FACT to the kinetochore requires intact CCAN (Fig. 1I-J), we wanted to investigate how individual CCAN subcomplexes contribute to FACT localization. RNA interference (RNAi) was used to deplete CCAN subcomplexes in RPE-1 cells and mitotic cells were immunostained for SSRP1. RPE-1 cells were treated with siRNA against CENP-HIKM for 72 hours to deplete the complex from the kinetochore (Fig. S5A-B,D). As a result, localization of SSRP1 was severely affected. CENP-OPQUR and CENP-TW localization was also substantially reduced, indicating that depletion of the CENP-HIKM subcomplex destabilizes CCAN (Fig. 3A-C, Fig. S5E,F). Conversely, CENP-A was not perturbed upon CENP-HIKM depletion (Fig. S5B,C).

**Figure 3.**
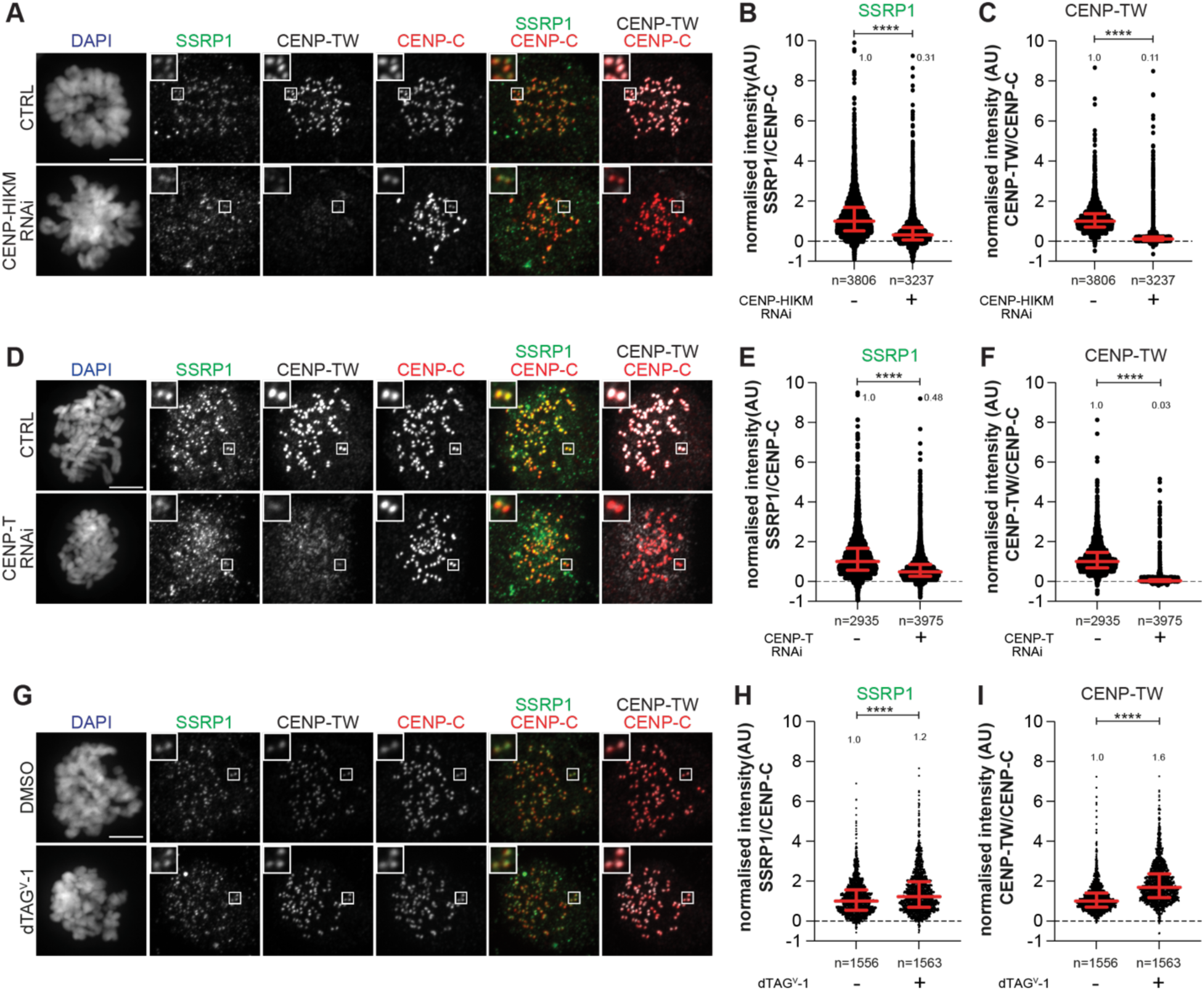
Mitotic localization of FACT depends on CENP-HIKM and -TW, but not on CENP-OPQUR. **(A)** Representative images of localization of SSRP1 and CENP-TW after depletion of CENP-HIKM complex in RPE-1 cells. CENP-HIKM RNAi was performed for 72 hours using silencing oligonucleotides for each subunit at 30 nM concentration. Cells were treated with STLC (5 µM) for 16 hours prior to fixation to obtain a mitotic population of cells. CENP-C identifies kinetochores and DAPI stains DNA. Three biological replicates were performed. Scale bar: 5 μm. (**B**) Scatter plots of SSRP1 levels at kinetochores for the experiment shown in panel (A). *n* refers to individually measured kinetochores. (**C**) Scatter plots show quantification of CENP-TW levels at kinetochores of the experiment shown in panel (A). *n* refers to individually measured kinetochores. (**D**) Representative images of localization of SSRP1 and CENP-TW after depletion of CENP-T complex in RPE-1 cells. CENP-T RNAi was performed for 60 hours using oligos for each subunit at 30 nM concentration. Cells were treated with Nocodazole (3.3 µM) for 4 hours prior to fixation to obtain mitotic population. CENP-C was used to visualize kinetochores and DAPI to stain DNA. Three biological replicates were performed. Scale bar: 5 μm. (**E**) Scatter plots show quantification of SSRP1 levels at kinetochores of the experiment shown in panel (D). *n* is the number of individually measured kinetochores. (**F**) Scatter plots show quantification of CENP-TW levels at kinetochores of the experiment shown in panel (D). *n* is the number of individually measured kinetochores. (**G**) Representative images of localization of SSRP1 and CENP-TW after depletion of CENP-U complex in RPE-1-CENP-U-FKBP^F36V^ cells. Cells were treated with dtag^V^-1(500 nM) for 24 hours to degrade endogenous CENP-U. Cells were treated with Nocodazole (3.3 µM) for 4 hours prior to fixation to obtain mitotic population. CENP-C identifies kinetochores and DAPI stains DNA. Three biological replicates were performed. Scale bar: 5 μm. (**H**) Scatter plots of SSRP1 levels at kinetochores for the experiment shown in panel (G). *n* is the number of individually measured kinetochores. (**I**) Scatter plots of CENP-TW levels at kinetochores for the experiment shown in panel (G). *n* is the number of individually measured kinetochores.

A 60-hour CENP-T RNAi treatment eliminated CENP-TW from the kinetochore (Fig. S5G, Fig. 3D,F). Also in this case, a concomitant decrease in the kinetochore levels of FACT, CENP-HK, and CENP-O was observed, whereas the levels of CENP-A remained stable (Fig. 3D,E; Fig. S5H-L). While rapid depletion of CENP-C showed that FACT localization does not directly depend on CENP-C (Fig. 1G-H), CENP-C was also not sufficient to retain FACT at the kinetochore (Fig. 3A-F). Thus, collectively, the CCAN subcomplexes are inter-dependent for their localization, in agreement with previous literature (*19*, *63*, *73–75*). Furthermore, our results demonstrate that FACT localization at the kinetochore during mitosis depends on the CCAN (Fig. 1I-J, Fig. 3A-F).

To assess a potential contribution of CENP-OPQUR to the recruitment of FACT, we endogenously tagged both alleles of CENP-U with FKBP^F26V^ and used the resulting cell line to rapidly degrade CENP-U through addition of dTAG^V^-1 (*76*). A 24-hour treatment led to the complete loss of CENP-U and CENP-R from the kinetochore, suggesting that the entire CENP-OPQUR complex, not only CENP-U, are removed. Furthermore, Polo-like kinase 1 (PLK1) localization, which partially depends on CENP-OPQUR (*75*), decreased (Fig. S6A-F). On the other hand, localization of CENP-TW and CENP-HK did not require CENP-U (Fig. 3G,I; Fig. S6G,I). In fact, CENP-TW displayed an increase in its kinetochore levels (Fig. 3I). This may indicate competition between CENP-TW and -OPQUR within the CCAN, but may also reflect a staining artifact caused by enhanced accessibility of the antigen. Finally, FACT localization was not affected by the depletion of CENP-U (Fig. 3G-H), indicating that CENP-OPQUR is not necessary for recruiting or retaining FACT at the kinetochore, even if it interacts with FACT *in vivo*, as suggested by co-immunoprecipitation (Fig. S4D). Alternatively, CENP-OPQUR and FACT may interact in a separate complex outside of CCAN. Thus, collectively, our results demonstrate the importance of CCAN, even if we cannot point to a single CCAN subunit as a recruiter of FACT. Kinetochore localization of FACT is substantially reduced upon depletion of CENP-HIKM or CENP-TW, a condition that additionally triggers a reduction of CCAN stability. Conversely, CENP-C and CENP-OPQUR are not strictly required for the localization of FACT to kinetochores.

### FACT dimerization and Mid-AID domains are required for CCAN binding

The FACT subunits SPT16 and SSRP1 are multi-domain distant paralogs with distinct functions (*77*). To identify binding sites for CCAN, we produced different truncations or isolated domains of FACT (Fig. 4A) and used them as preys in pull-down assays with CCAN subcomplexes as baits. As already observed, ^MBP^CENP-C^EGFP^ and FACT bound directly (Fig. S4B, Fig. 4B). Additionally, ^MBP^CENP-C^EGFP^ pulled down SPT16^Mid-AID^, a minimal SPT16, fragment. It also pulled down, with apparently slightly higher affinity, construct 2 (SPT16^508-988^/SSRP1^1-514^, henceforth FACT^trunc^) (Fig. 4A,B). GST-tagged CENP-OPQUR interacted robustly with FACT and FACT^trunc^, but only weakly with SPT16^Mid-AID^ (Fig. 4C). Conversely, MBP-tagged CENP-TW (^MBP^CENP-TW) was sufficient to bind SPT16^Mid-AID^ (Fig. 4D). ^MBP^CENP-TW also bound strongly to FACT^trunc^ and with low affinity to SSRP1^Mid-AID^ (Fig. 4D). Thus, association of ^MBP^CENP-TW with either SPT16 and SSRP1 Mid-domain depended on the presence of an intact AID domain (Fig. 4D, lanes 3-6). In summary, FACT^trunc^ was sufficient to bind all CCAN subcomplexes, while the N-terminal aminopeptidase-like domain of SPT16 and the C-terminal HMG-domain of SSRP1 are dispensable for CCAN-binding (Fig. 4B-D). In a reverse pull-down, the apparent strengths of the interaction of CCAN with either full-length ^MBP^FACT or ^MBP^FACT^trunc^ were identical (Fig. 4E).

**Figure 4.**
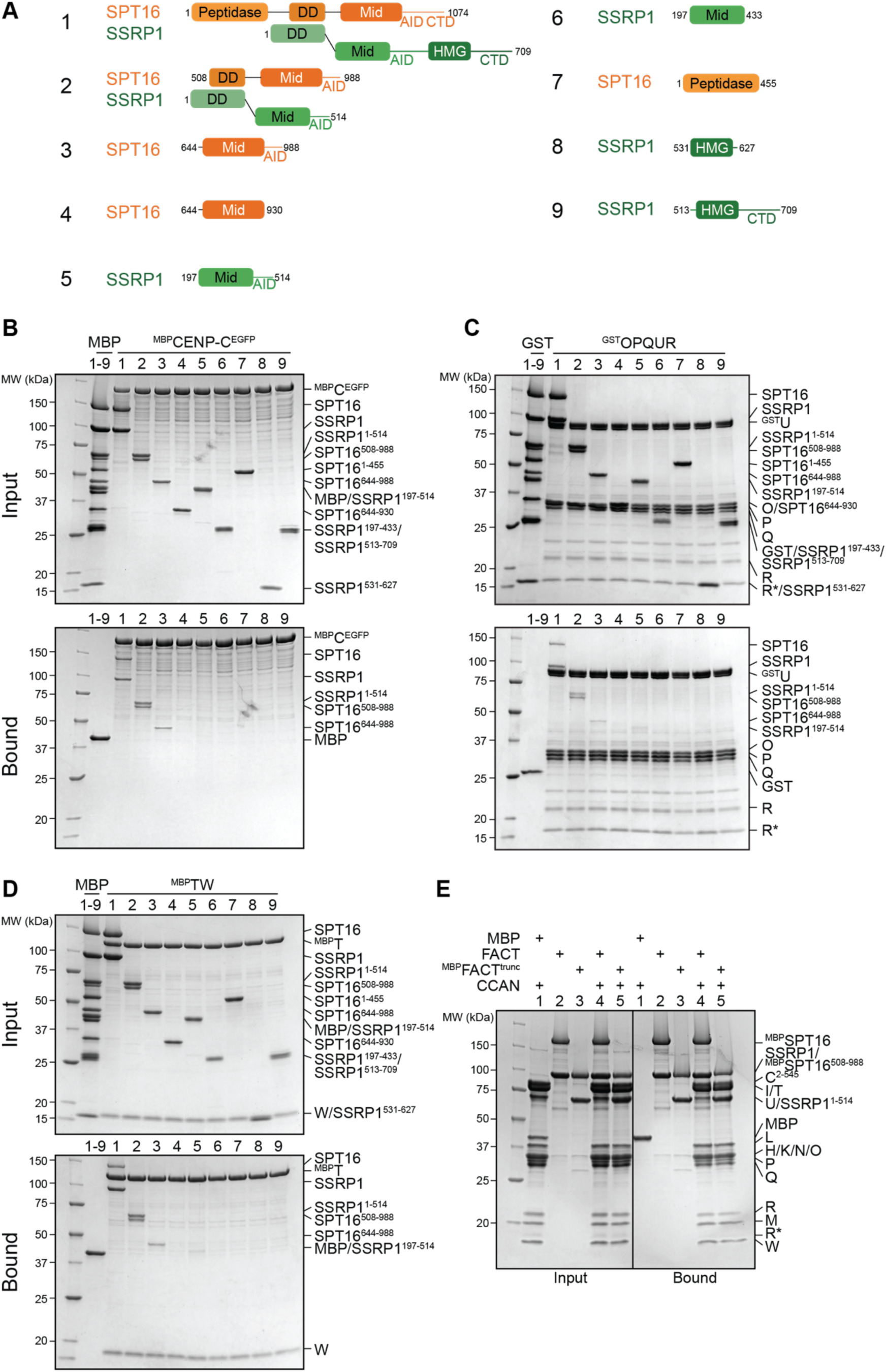
CCAN binding requires FACT dimerization and Mid-AID domains. (**A**) Scheme of FACT domains or truncations used as prey in the following pull-down assays. (**B**) Amylose-resin pull-down assay with ^MBP^CENP-C^EGFP^ to analyze binding of the FACT constructs in (A). (**C**) Glutathione-agarose pull down assay using CENP-OPQUR with CENP-U fused to an N-terminal GST as bait and FACT constructs in (A) as preys. (**D**) Amylose-resin pull-down assay with ^MBP^CENP-T/CENP-W as bait and FACT constructs in (A) as preys. (**E**) Amylose-resin pull-down assay comparing CCAN binding to ^MBP^FACT and ^MBP^FACT^trunc^.

### FACT requires phosphorylation by CK2 to interact with CCAN

FACT is regulated by, and also directly binds to, acidophilic casein kinase II (CK2) (*38*, *78–80*). The CK2 holoenzyme is a tetramer composed of the active subunit CK2α or CK2α’ and the regulatory and dimerizing subunit CK2β (*81*). CK2 is a promiscuous kinase with hundreds of different substrates involved in numerous biological processes and diseases (*82*). It is characterized as a constitutively active kinase and its regulation is not defined by a single mechanism, but rather is substrate specific (*83*). Despite its localization to different cellular compartments, CK2 is mostly active in the nucleus (*84*, *85*), where it has a role in transcription (*86*, *87*).

Recombinant FACT purified from insect cells was strongly phosphorylated, but treatment with λ-phosphatase removed phosphorylation (Fig. S7A). The elution volume of FACT was unaffected by changes in its phosphorylation status (Fig. S7B). Unexpectedly, dephosphorylated FACT (repurified to eliminate λ-phosphatase) failed to bind CCAN in a SEC co-elution assay (Fig. 5A). Addition of CK2 to the reaction to induce phosphorylation of FACT restored the binding of FACT/CCAN in analytical SEC (Fig. 5A). The phosphorylation dependency of FACT/CCAN complex formation was corroborated in a solid-phase assay (Fig. 5B, lanes 11-13). This assay was also used to probe the phosphorylation dependency of the interaction of FACT with specific CCAN subcomplexes. Dephosphorylated ^MBP^FACT failed to pull down CENP-C^2-545^HIKM, CENP-OPQUR and CENP-TW. These interactions were partially restored upon CK2 phosphorylation, although not to the levels observed with the sample before dephosphorylation (Fig. 5B), probably due to incomplete rephosphorylation (Fig. S7C). Some CCAN subunits, including CENP-C and/or CENP-I, CENP-U and CENP-T, were also phosphorylated by CK2 (Fig. S7C). Of note, additional kinases demonstrated an ability to phosphorylate FACT, but they could not restore the interaction with CCAN, indicating that the effects on CCAN binding are specific to CK2 (Fig. S7D).

**Figure 5.**
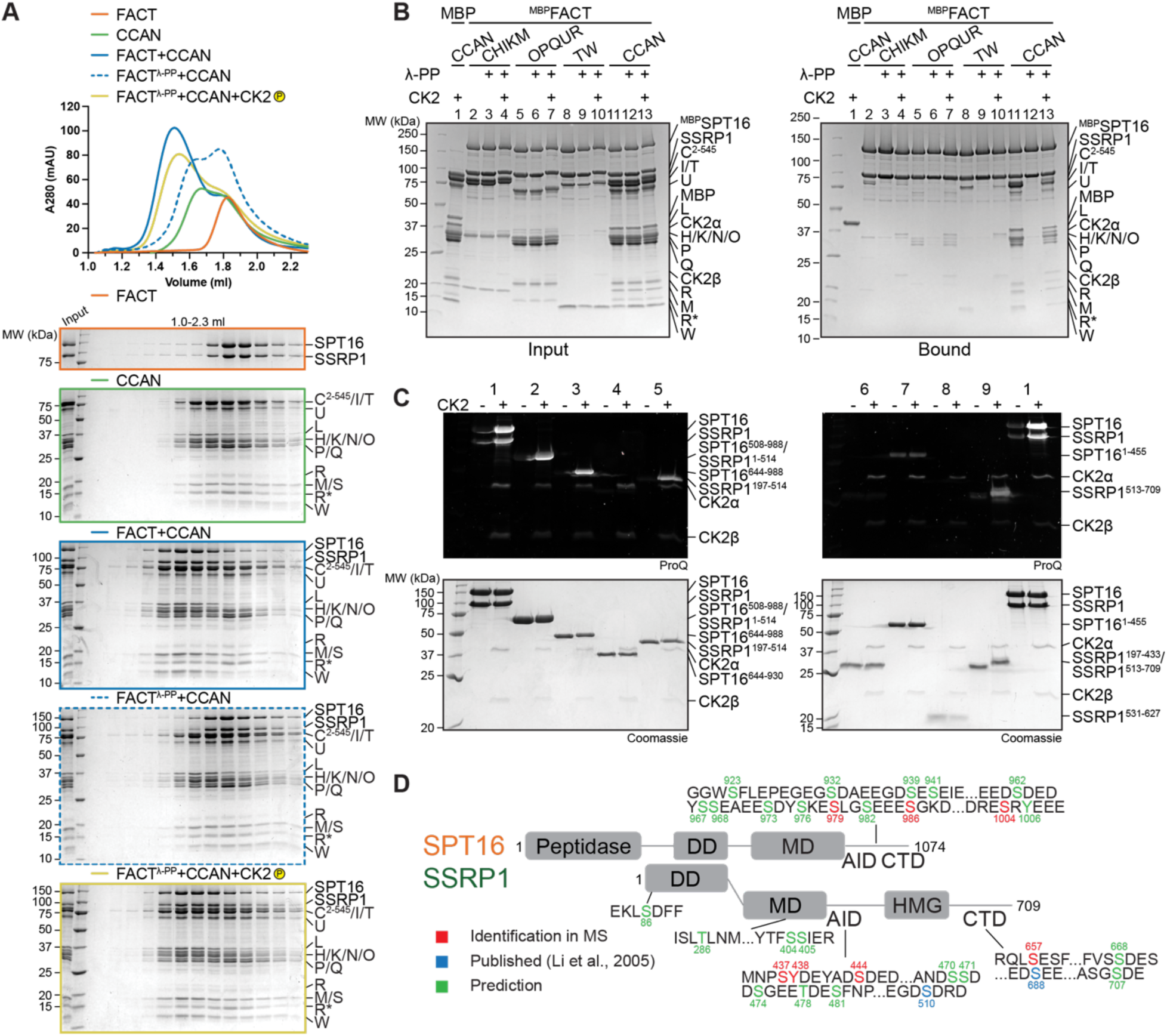
The FACT/CCAN interaction requires phosphorylation of FACT by CK2. (**A**) Analytical SEC comparing CCAN binding of untreated FACT, dephosphorylated, re-purified FACT and dephosphorylated FACT, where the sample is treated with CK2. The experiment was performed with samples in Figure 1D. The first three samples are the same as Figure 1D. (**B**) Amylose-resin pull-down assay using either untreated ^MBP^FACT or dephosphorylated ^MBP^FACT as bait and CCAN subcomplexes or CCAN as preys. λ-PP indicates that ^MBP^FACT had been dephosphorylated and λ-PP had been removed by SEC before the experiment. CK2 indicates that the sample was treated with CK2. (**C**) FACT constructs (Fig. 4A) were treated with CK2 and the phosphorylation state was assessed by ProQ Diamond staining. Coomassie staining of the corresponding SDS-PAGE gels are shown below. (**D**) Scheme of FACT with sequences of potential CK2 phosphorylation sites indicated. CK2 sites were based on mass spectrometry, predictions or previous publications.

FACT constructs described in Figure 4 were used to analyse which regions of the complex are phosphorylated by CK2. SPT16 and SSRP1 Mid-AID domains were phosphorylated in an AID-dependent manner (Fig. 5C), in agreement with the ability of CK2 to phosphorylate acidic sequences (*88*). Additionally, the C-terminal region of SSRP1 was also phosphorylated by CK2 (Fig. 5C). Phosphorylated FACT complex was subjected to mass spectrometry analysis to identify target sites. This analysis failed to identify the precise phosphorylation site within the AID sequences, but in combination with sequence-based prediction (*89*) and published phosphorylation sites (*79*), we propose more than thirty potential CK2 phosphorylation sites on FACT (Fig. 5D, Table S1). In conclusion, by dephosphorylating insect cell-expressed FACT, we revealed that the interaction with CCAN is phosphorylation-dependent. We identified the required kinase as CK2, but we could not determine specific phosphorylation sites necessary for binding. Due to the constitutive activity of CK2 (*83*), cellular conditions leading to phosphorylation of FACT by CK2 remain unclear.

### DNA competes with FACT for CCAN binding

CCAN decodes CENP-A^NCP^ through CENP-C and CENP-N and binds DNA cooperatively (*9–15*, *26*), while FACT only binds to nucleosomes that are partially destabilized, e.g. by an actively transcribing RNA polymerase II (*55*, *90–93*). We set out to further dissect FACT’s interaction with CCAN on chromatin using biochemical reconstitution. In analytical SEC, addition of a 145-bp DNA fragment prevented assembly of the FACT/CCAN complex altogether, as DNA binding to CCAN displaced FACT from the complex (Fig. 6A). The same effect was observed upon addition of a 75-bp DNA fragment (Fig. S8A). This result was confirmed in a solid-phase assay (Fig. 6B, lanes 8,9). CCAN binds DNA very tightly, whereas individual subcomplexes bind to DNA with much lower affinity, if at all (*26*, *27*). We therefore asked how DNA influenced the interaction of FACT with the CCAN subcomplexes CENP-C^2-545^HIKM, CENP-OPQUR and CENP-TW. Although a marginal reduction in binding of each complex was detected (Fig. 6B, lanes 2-7), the effect of DNA was considerably less pronounced than in presence of the complete CCAN (Fig. 6B).

**Figure 6.**
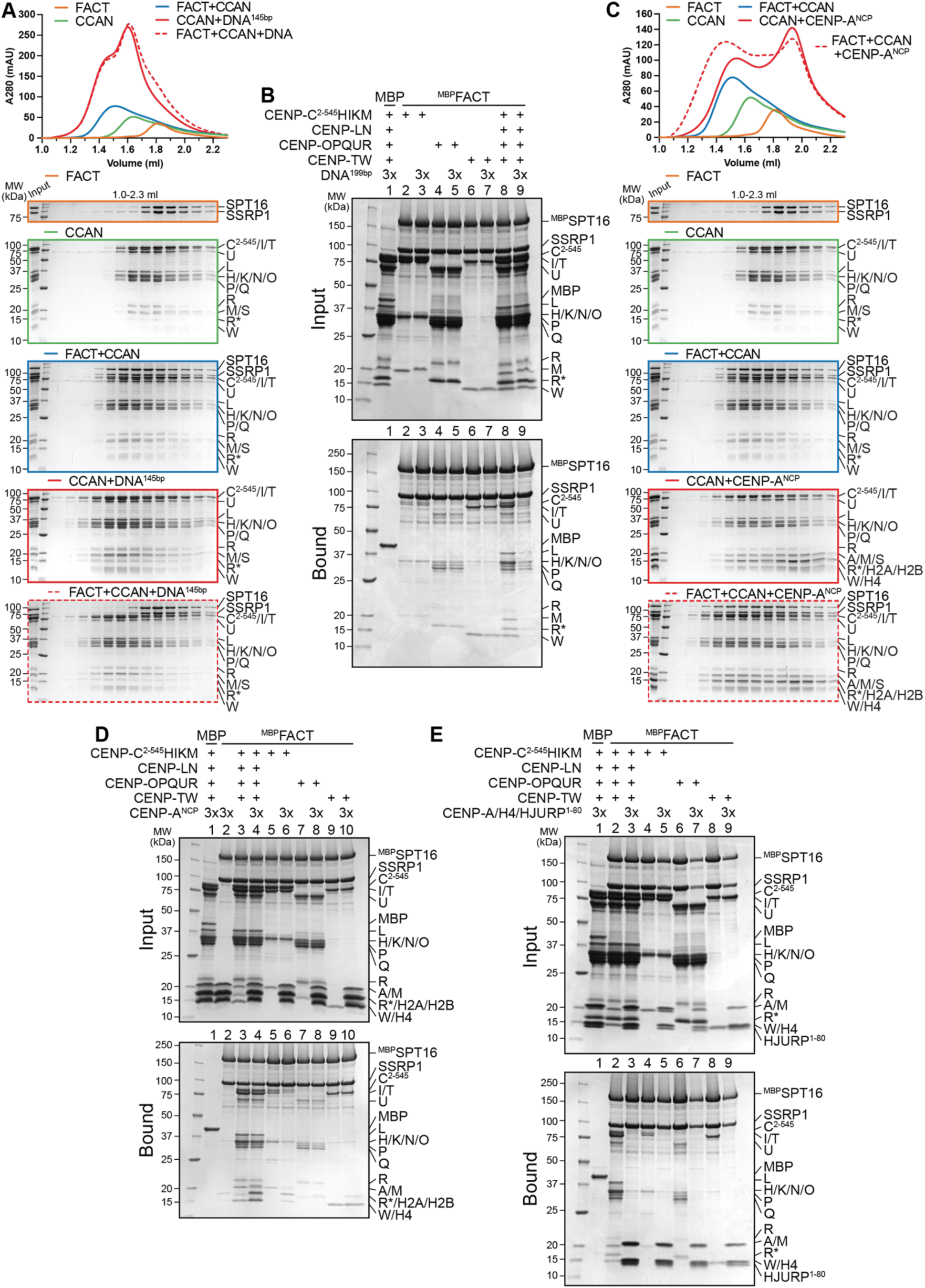
FACT competes with DNA for CCAN binding and CCAN competes with centromeric histones for FACT binding. (**A**) Analytical SEC to test the effect of DNA on the FACT/CCAN complex. A 145-bp Widom 601 sequence was used. (**B**) Amylose-resin pull-down assay with ^MBP^FACT as bait and CCAN complexes or full CCAN and DNA as preys. A 199-bp centromere 1 (CEN1)-like DNA sequence was used. (**C**) Analytical SEC to assess reconstitution of a FACT/CCAN/CENP-A^NCP^ complex. The experiment is part of a larger experiment that includes the experiment in panel (A). The CENP-A/H4/H2A/H2B histone octamer was reconstituted on 145-bp Widom 601 sequence. The SDS-PAGE gels of FACT, CCAN, and FACT+CCAN are duplicates of those shown in panel (A). The chromatograms are also the same but displayed on a different scale. (**D**) Amylose-resin pull-down assay of ^MBP^FACT and CCAN subcomplexes or CCAN with the addition of an excess of CENP-A^NCP^. (**E**) Amylose-resin pull-down assay of ^MBP^FACT and CCAN subcomplexes or CCAN with the addition of an excess of CENP-A/H4/HJURP^1-80^.

Next, we tested whether CCAN binding to FACT is compatible with binding to a CENP-A nucleosome core particle (NCP, on a 145 bp Widom 601 sequence). Recombinant FACT did not bind to DNA or intact nucleosomes in analytical SEC (Fig. S8B). When mixed with CCAN and a CENP-A^NCP^, however, a tripartite complex formed (Fig. 6C). These observations were corroborated using a pull-down assay, where CENP-A^NCP^ was seen to interact with ^MBP^FACT through CCAN (Fig. 6D). The association of CCAN subcomplexes was also evaluated. CENP-OPQUR and CENP-TW did not bind to nucleosomes and the interaction with FACT was essentially unaltered. Binding of CENP-C^2-545^HIKM was marginally reduced upon the addition of CENP-A^NCP^ (Fig. 6D). We ligated a naked DNA sequence to a CENP-A nucleosome built on α-satellite DNA, creating a 348-bp sequence of which roughly half was embedded in a nucleosome. The overhanging DNA acted comparably to free DNA, effectively displacing FACT from a CCAN/CENP-A^NCP^ complex (Fig. S8C,D). Collectively, these results suggest that FACT may recognize a form of CCAN that is not directly bound to DNA at centromeres, despite substantial biochemical and structural information indicating that CCAN binds DNA tightly through the CENP-LN vault and potentially through various neighbouring DNA-binding structures (*26–28*, *94*). Whether a form of CCAN devoid of DNA is present at kinetochores during mitosis, however, remains unclear.

### FACT cannot bind centromeric histones and CCAN simultaneously

As a histone chaperone, FACT is known to bind to both H2A/H2B dimers and H3/H4 tetramers (*54*, *55*). H2A/H2B and CENP-TW compete for the same binding site on FACT, and FACT has a binding preference for the former (*62*). We wanted to broaden our analysis of the FACT/CCAN complex in relation to centromeric histones. A trimeric complex of histones CENP-A/H4 with the first 80 residues of its chaperone HJURP was added to a pull-down assay with ^MBP^FACT as bait and CCAN as prey. CENP-A/H4/HJURP^1-80^ outcompeted CCAN in FACT binding (Fig. 6E). This result was confirmed in an orthogonal assay, where ^MBP^HJURP^1-80^ in complex with CENP-A/H4 as bait bound FACT efficiently, but CCAN was excluded from the complex (Fig. S9A). ^MBP^HJURP^1-80^ in absence of CENP-A/H4, used as control, did not bind FACT nor CCAN (Fig. S9A), indicating that competition with CCAN for FACT binding is caused by the CENP-A/H4 dimer rather than the chaperone. Binding of CCAN subcomplexes to FACT was also disrupted upon addition of CENP-A/H4/HJURP^1-80^ (Fig. 6E), suggesting that CENP-A/H4 and CCAN may share the same binding site on FACT. This was validated by testing the binding of different FACT constructs to CENP-A/H4/^MBP^HJURP^1-80^. Indeed, Mid-domains and AID segments of SPT16, but also SSRP1, were important for binding to centromeric histones (Fig. S9B). These data suggest that FACT is not able to chaperone CENP-A/H4 and CCAN simultaneously.

## Discussion

FACT has established roles in transcription, replication, and DNA repair. On the other hand, the functional significance of its enrichment at human kinetochores remains unclear. We identified novel interactions of FACT with CCAN subunits CENP-C and CENP-OPQUR in addition to the previously reported binding to the HFD-containing complex CENP-TW (*62*). These interactions act cooperatively to form a stable 18-subunit complex. During mitosis, FACT localizes to the kinetochore in a CCAN-dependent manner, and the CENP-HIKM and CENP-TW complexes are especially important for this localization. The direct and specific interaction of FACT with the kinetochore suggests a role linked to kinetochore assembly and the regulation of kinetochore interactions with centromeric chromatin. The exact function of kinetochore FACT will have to be elucidated, but our study paves the way for a detailed functional analysis.

*In vitro*, FACT was displaced from the CCAN by DNA, suggesting that CCAN is not stably anchored to chromatin while FACT is bound. This is unexpected, as FACT localizes to the kinetochore in a CCAN-dependent manner during mitosis (Fig. 1I-J), a time when we expect a tight connection between kinetochores and centromeric DNA. Our results may suggest that a subset of CCANs complexes is engaged with FACT rather than with DNA. There is only limited information on the mechanism and timing of recruitment of CCAN to the centromere during or after DNA replication. In line with its chaperone activity, FACT may stabilize the CCAN or CCAN subunits in solution and help in the deposition and assembly of the CCAN at the centromere. We observed that while individual interaction partners had a lower affinity for FACT than the entire CCAN, CENP-C^2-545^HIKMLNTW^HFD^ was the strongest binder, while CENP-TW and -OPQUR were moderately anti-cooperative and competing for FACT binding (Fig. 2A-E, H).

Our observations also indicate that individual interactions of FACT and CCAN ultimately cooperate to enhance overall binding. For instance, we suspect that the interaction of FACT with CENP-OPQUR may undergo a rearrangement inside the CCAN in comparison to the isolated FACT/CENP-OPQUR complex. Thus, FACT may preferentially bind to CCAN complexes that are not fully assembled or properly incorporated into centromeric chromatin. This possibility may also partly explain some discrepancies between the results of binding assays *in vitro* and the analysis of FACT localization after displacement of CCAN subunits *in vivo*. For instance, the CENP-HIKM complex appeared to have a disproportionate effect on FACT localization if gauged against the apparently low binding affinity for FACT *in vitro*. Given the position of CENP-HIKM in the CCAN hierarchy, which is upstream compared to other sub-complexes, it is reasonable to assume that its depletion would lead to a more significant effect on FACT recruitment to the kinetochore. We surmise that depletion of CENP-HIKM may indirectly affect the interaction of CENP-TW with DNA, causing FACT displacement indirectly.

Upon re-entry into interphase, FACT redistributes to the entire chromosome, more pronouncedly around nucleoli (*65*, *66*) (Fig. 1F). Our inability to visualize FACT at kinetochores outside of mitosis does not necessarily imply depletion of FACT from these structures, as visualizing kinetochore FACT by immunofluorescence during interphase against a more pronounced chromosome signal is technically challenging. As CCAN localizes to the centromere throughout the cell cycle (*6*), it will be important to establish if FACT acts there outside of mitosis. Chromatin is considered transcriptionally silent in mitosis (*95*), but it has been suggested that centromeric transcription is also active during mitosis (*69–72*). Thus, it is possible that the localization of FACT at the kinetochore coincides with active centromeric transcription in mitosis and interphase. For instance, FACT may prevent loss of CCAN as RNA polymerase II passes through centromeric chromatin. This would be reminiscent of FACT’s known role in preventing histone loss during transcription (*36*, *96*).

Due to FACT’s role in multiple chromatin related mechanisms, studying its specific role at the kinetochore is challenging. As RNAi-based depletion of FACT is slow and likely causative of pleiotropic effects, targeted acute degradation of FACT specifically in mitosis will be required to overcome this challenge and focus on its mitosis-specific role. Another option would be to identify separation-of-function mutant to target specific functions of FACT. The AID of SPT16 as well as phosphorylation by CK2 are important for other functions in addition to mediating the interaction with CCAN and are therefore not appropriate targets for mutations. Investigation of potential CK2 sites on FACT may ultimately identify sites that are solely important for CCAN binding. Finally, structural information on a FACT/CCAN complex could facilitate the identification of specific interaction interfaces. So far, our efforts to obtain high-resolution structures of the CCAN/FACT complex have been thwarted by lack of order of the resulting complexes.

The phosphorylation of FACT by CK2 is indispensable for FACT/CCAN complex formation. *In vitro*, binding of DNA to CCAN leads to the dissociation of FACT, while FACT preferentially binds to centromeric histones, which share binding sites with CCAN. Collectively, these data suggest that FACT impacts the kinetochore directly rather than sharing the same function at the centromere as in other parts of chromatin. FACT is predicted to possess up to thirty or more CK2 phosphorylation sites, especially in the AID sequences (Fig. 5D). Currently, it remains uncertain which of these sites are crucial for the binding to CCAN. Nevertheless, phosphorylation of FACT is also important for other functions (*38*, *78–80*) and it is not clear whether the different CK2 sites on FACT are functionally related. Interestingly, phosphorylation of FACT reduces its DNA binding activity (*79*, *97*). It is possible that FACT changes its exact localization from DNA to histones or the CCAN depending on its phosphorylation state. Alternatively, different pools of FACT may accomplish different functions simultaneously. For instance, one pool may bind CCAN, while another may bind CENP-A/H4 or other histones during transcription and replication. In summary, we provided a characterization of the FACT/CCAN interaction *in vitro* and *in vivo*, and set the basis for future work aiming to dissect this interaction.

## Materials and Methods

### Plasmids

Plasmids for the expression of CENP-C^2-545^HIKM, CENP-C^2-545^, CENP-C^189-400^, ^MBP^CENP-C^721-C^, CENP-HIKM, CENP-LN, CENP-OPQUR, CENP-OPQ^68-C^U^115-C^R CENP-TWSX, CENP-TW, CENP-T^458-C^W, CENP-SX, ^MBP^CENP-T/W, CENP-A/H4, H2A/H2B, CDK1, Cyclin B, CKS1, PLK1, Aurora B^45-344^/INCENP^835-903^ and for the production of DNA sequences were generated as previously described (*12*, *13*, *20*, *26*, *63*, *64*, *74*, *75*, *98*, *99*). Plasmid expressing human CK2α^1-335^ and CK2β^1-193^ were a kind gift of K. Niefind (University of Cologne, Germany). SPT16 and SSRP1 with a N-terminal His-tag and a TEV cleavage site were cloned into a pFL-derived MultiBac vector (*100*). Sequences of ^MBP-TEV^SPT16, ^His-TEV^SSRP1, ^His-TEV^SPT16^644-988^ (Mid-AID), ^His-TEV^SPT16^644-930^ (Mid), ^His-TEV^SSRP1^197-514^ (Mid-AID), ^His-TEV^SSRP1^197-433^ (Mid), ^His-TEV^SPT16^508-988^, ^MBP-TEV^SPT16^508-988^, SSRP1^1-514^, ^His-TEV^SSRP1^1-514^, ^His-MBP^CENP-C^EGFP^ and ^GST^CENP-U were inserted into pLIB vectors. These were used to combine ^MBP-TEV^SPT16+^His-TEV^SSRP1 (^MBP^FACT), ^His-TEV^SPT16^508-988^+SSRP1^1-514^ (FACT^trunc^), ^MBP-TEV^SPT16^508-988^+ ^His-TEV^SSRP1^1-514^ (^MBP^FACT^trunc^), ^GST^CENP-U with previously described CENP-O/P/Q/R and ^His-TEV^CENP-Q^68-C^ and CENP-U^115-C^ with CENP-O/P/R (*63*) in pBIG1a vectors for baculovirus based multigene-expression (*101*). ^His-PreSc^SPT16^1-544^ (Peptidase-like domain), ^His-PreSc^SSRP1^531-627^ (HMG domain), ^His-PreSc^SSRP1^513-C^, ^MBP^CENP-C^2-400-His^, ^MBP^CENP-C^401-600-His^, ^MBP^CENP-C^401-545-His^, ^MBP^CENP-C^546-600-His^, ^MBP^CENP-C^601-720-His^, ^MBP^CENP-C^721-759-His^, ^His-MBP-TEV^CENP-C^760-C^, ^MBP-TEV^HJURP^1-80-His^ and ^His-PreSc^CENP-A co-expressed with H4 and ^MBP-TEV^HJURP^1-80^ were cloned into a pETDuet vector using Gibson cloning (*102*).

### Purification of DNA fragments

Generation of the Widom 601 145-bp (ATCAGAATCCCGGTGCCGAGGCCGCTCAATTG GTCGTAGACAGCTCTAGCACCGCTTAAACGCACGTACGCGCTGTCCCCCGCGTTTTAA CCGCCAAGGGGATTACTCCCTAGTCTCCAGGCACGTGTCAGATATATACATCGAT) and the CEN1 (centromere 1)-like 75-bp (ATCCGTGGTAGAATAGGAAAT ATCTTCCTATAGAAACTAGACAGAATGATTCTCAGAAACTCCTTTGTGATGGAT), 165-bp (GTGGTAGAATAGGAAATATCTTCCTATAGAAACTAGACAGAATGATTCT CAGAAACTCCTTTGTGATGTGTGCGTTCAACTCACAGAGTTTAACCTTTCTTTTCATAG AGCAGTTAGGAAACACTCTGTTTGTAATGTCTGCAAGTGGATATTCAGACGCCCTTG, 183-bp (AGGCCTTCGTTGGAAACGGGATTTCTTCATATTCTGCTAGACAGAAGAATTCTCAGTAACTTCCTTGTGTTGTGTGTATTCAACTCACAGAGTTGAACG ATCCTTTACACAGAGCAGACTTGAAACACTCTTTTTGTGGAATTTGCAGGCCTAGATTTCAGCCGCTTTGAGGTCAATCACCCC) and 199-bp (ATCGCCCTTGAG GCCTTCGTTGGAAACGGGATTTCTTCATATTCTGCTAGACAGAAGAATTCTCAGTAACTTCCTTGTGTTGTGTGTATTCAACTCACAGAGTTGAACGATCCTTTACACAGAGCAGA CTTGAAACACTCTTTTTGTGGAATTTGCAGGCCTAGATTTCAGCCGCTTTGAGGTCAATCACCCCGTGGAT) was performed as previously described (*20*, *26*).

### Reconstitution of nucleosomes

CENP-A nucleosome core particles on 145-bp 601 or CEN1-like 183-bp DNA were produced as previously reported (*103*).

To generate CENP-A nucleosomes on 348 bp DNA, 165-bp DNA was ligated to the front of 183-bp DNA on pre-reconstituted CENP-A nucleosome. The two species were mixed in equimolar amounts based on the concentration of the DNA fragements in a buffer consisting of 10 mM Tris pH 7.4, 100 mM NaCl and 1 mM EDTA. Two times the amount of ^MBP^T4 DNA ligase^His^ (produced in house) relative to the DNA fragment was added with 10X T4 DNA Ligase Buffer and the reaction was incubated for approximately 16 h at 4 °C. The reaction was passed through two consecutive 1 ml HisTrap FF columns (Cytiva), equilibrated in 10 mM Tris pH 7.4 and 100 mM NaCl to remove His-tagged T4 DNA ligase. The flow-through and wash was collected and EDTA was added to a final concentration of 2 mM.

### Protein expression and purification

CENP-C^2-545^HIKM, CENP-C^2-545^, CENP-C^189-400^, CENP-HIKM, CENP-LN, CENP-OPQUR, CENP-TWSX, CENP-TW, CENP-T^458-C^W, CENP-SX, ^MBP^CENP-T/W, CDK1/Cyclin B/CKS1, PLK1 and Aurora B^45-344^/INCENP^835-903^ were expressed and purified as previously reported (*12*, *13*, *20*, *26*, *63*, *64*, *74*, *75*, *98*, *99*). ^GST^CENP-OPQUR and CENP-OPQ^68-C^U^115-C^R were purified identically to the wild type (*63*) by using either glutathione affinity or Nickel affinity as a first step.

FACT, ^MBP^FACT, FACT^trunc^, ^MBP^FACT^trunc^, SPT16^Mid-AID^, Mid domain, SSRP1^Mid-AID^ and Mid domain were expressed by infecting Tnao38 cells with a virus:culture ratio of 1:20 and incubating the cells at 27 °C for 72 h. For the expression of ^MBP^CENP-C^EGFP^ a virus:culture ratio of 1:40 was used. SPT16 peptidase-like domain, SSRP1^HMG^, SSRP1^513-709^, ^MBP^HJURP^1-80-His^ and ^His^CENP-A/H4/^MBP^HJURP^1-80^ were expressed in *E. coli* BL21(DE3)-Codon-plus-RIL cells by growing transformed cells to an OD_600_ of 0.7 in TB medium supplemented with ampicillin and chloramphenicol at 25 °C. Expression was induced by adding 0.1 mM IPTG and cells were cultured for 16 hours at 18 °C.

All purification steps were performed at 4 °C or samples were kept on ice. If not otherwise indicated, cells were resuspended in lysis buffer supplemented with Protease-Inhibitor Mix HP Plus (Serva), 1 mM PMSF and 10 µg/ml DNaseI and lysed by sonication. The lysate was subsequently clarified by centrifugation for 45 min at 10,0000 xg at 4 °C and filtration. After the final purification step, proteins of interest (POI) were concentrated, flash-frozen in liquid nitrogen and stored at -72 or -80 °C.

Cells expressing FACT, ^MBP^FACT, FACT^trunc^, ^MBP^FACT^trunc^, SPT16^Mid-AID^, Mid domain, SSRP1^Mid-AID^, Mid domain and SPT16 peptidase-like domain were resuspended in a buffer containing 20 mM Tris-HCl pH 8.0, 300 mM NaCl, 5% glycerol and 1 mM TCEP (Buffer A). The lysate was applied to a 5 ml HisTrap FF (Cytiva). The column was first extensively washed with Buffer A and then with Buffer A including 30 mM imidazole. Full-length FACT was eluted by a linear gradient to 400 mM imidazole, others were eluted in Buffer A with 250 mM imidazole. The fractions containing protein were pooled and diluted 1:4 with Buffer A containing 150 mM NaCl. This was loaded on two sequential 1 ml HiTrap Q HP (Cytiva) anion exchange columns. The columns were washed and protein was eluted by a gradient to 1 M NaCl. Peak fractions were analyzed in SDS-PAGE and fractions containing protein or a stoichiometric complex were pooled and concentrated. To obtain dephosphorylated protein FACT and ^MBP^FACT were treated with λ-phosphatase (produced in house) at 4 °C in presence of 1 mM MnCl_2_ for approximately 16 h. Full-length FACT, ^MBP^FACT, FACT^trunc^ and ^MBP^FACT^trunc^ were finally applied to a HiLoad 16/600 Superose 6 pg column, the others were purified on a HiLoad 16/600 Superdex 200 pg (Cytiva).

Expressions of SSRP1^HMG^ and SSRP1^513-709^ were resuspended in a buffer composed of 20 mM HEPES pH 6.8, 300 mM NaCl, 5% glycerol, 10 mM imidazole and 1 mM TCEP. Nickel affinity purification was performed as explained above. Protein was diluted 1:4 in the same buffer with 100 mM NaCl and loaded on a 5 ml HiTrap Heparin HP column (Cytiva) and protein was eluted in a gradient to 1 M NaCl. Fractions containing the protein were pooled and concentrated to be applied to a HiLoad 16/600 Superdex 75 pg (Cytiva).

A pellet of ^MBP^CENP-C^EGFP^ expressing Tnap38 cells was resuspended in approximately 10 volumes of TALON buffer (50 mM Hepes pH 7.0, 500 mM NaCl, 5 mM MgCl_2_, 5% glycerol, 5 mM imidazole, 2 mM TCEP) supplemented with 2 mM PMSF and DNaseI. Affinity purification was performed on a 5 ml HisTALON Cartridge pre-packed with TALON Superflow Resin (Cytiva) and the column was washed with 10 CV buffer. The protein was eluted in TALON buffer A with 200 mM imidazole and subsequently diluted to 300 mM NaCl in Heparin buffer (20 mM Hepes pH 7.0, 5% glycerol, 2 mM TCEP). A 5 ml HiTrap Heparin HP column (Cytiva) was equilibrated in Heparin buffer including 300 mM NaCl and the diluted protein was bound to it. The column was washed with Heparin buffer with 300 mM NaCl and the protein was eluted in a linear gradient to 1 M NaCl in 150 ml. Peak fractions containing the POI were concentrated and subjected to SEC on a HiLoad 16/600 Superose 6 pg (Cytiva) in SEC buffer (20 mM Hepes pH 7.0, 500 mM NaCl, 5% glycerol, 1 mM TCEP^MBP^CENP-C^2-400^ and ^MBP^CENP-C^721-C^ were purified in a buffer consisting of 50 mM Hepes, pH 7.5, 500 mM NaCl, 10% glycerol and 1 mM TCEP. Proteins were purified on a 5 ml HisTrap FF (Cytiva) as indicated above and eluted in 250 mM imidazole. The eluate was diluted 3 times in Heparin buffer (20 mM Hepes pH 7.5, 150 mM NaCl, 5% glycerol, 1 mM TCEP), bound to a 5 ml HiTrap Heparin HP column (Cytiva) and eluted by a linear gradient to 1 M NaCl. Subsequently to SDS-PAGE, relevant fractions of ^MBP^CENP-C^2-400^ were further purified by SEC on a HiLoad 16/600 Superdex 200 pg, while ^MBP^CENP-C^721-C^, which dimerizes, was applied to a HiLoad 16/600 Superose 6 pg (Cytiva). ^MBP^CENP-C^401-600^, ^MBP^CENP-C^401-545^, ^MBP^CENP-C^546-600^, ^MBP^CENP-C^721-759^ and ^MBP^CENP-C^760-C^ were obtained by Nickel affinity purification in 20 mM Hepes pH 7.5, 500 mM NaCl, 10% glycerol, 10 mM imidazole and 1 mM TCEP. Proteins were eluted in 250 mM imidazole, concentrated and applied to a HiLoad 16/600 Superdex 200 pg (Cytiva) in 20 mM Hepes pH 7.5, 300 mM NaCl, 5% glycerol and 1 mM TCEP.

^MBP^CENP-C^601-720^ was purified on a 5 ml HisTrap FF (Cytiva) in 20 mM Tris pH 8.0, 300 mM NaCl, 5% glycerol and 1 mM TCEP and eluted in 250 mM imidazole. The eluate was diluted 5 times in 20 mM Tris pH 8.0, 200 mM NaCl, 5% glycerol and 1 mM TCEP and applied to two consecutive 1 ml HiTrap Q HP (Cytiva) columns. The POI was collected in the flow-through, while DNA bound to the column. The flow-through was concentrated and purified on a HiLoad 16/600 Superdex 200 pg (Cytiva)

Cells expressing ^His^CENP-A/H4/^MBP^HJURP^1-80^ were resuspended in a buffer consisting of 20 mM Tris pH 8.0, 1 M NaCl and 1 mM TCEP. The lysate was applied to a 5 ml HisTrap FF (Cytiva) column, which was first washed with buffer and subsequently with buffer including 10 mM imidazole. The protein was eluted in 250 mM imidazole and diluted 1:4 with IEX buffer (20 mM HEPES pH 6.8, 600 mM NaCl, 1 mM TCEP) and loaded on a 1 ml HiTrap SP HP (Cytiva). The protein was eluted by a linear gradient to 2 M NaCl and afterwards concentrated and purified on a HiLoad 16/600 Superdex 200 pg (Cytiva) in the initial buffer including 1 M NaCl. To obtain untagged protein, the eluate of Nickel affinity purification was treated with PreScission and TEV protease for approximately 16 h at 4 °C while it was dialyzed to a buffer containing 750 mM NaCl. Purification on a 1 ml HiTrap SP HP and SEC on a HiLoad 16/600 Superdex 200 pg (Cytiva) was performed as explained above. ^MBP^HJURP^1-80^ expressing E. coli were resuspended in a buffer consisting of 20 mM Hepes pH 7.5, 300 mM NaCl, 5% glycerol and 1 mM TCEP. The cells were lysed by high pressure in a microfluidizer and afterwards clarified by centrifugation. Nickel affinity purification was performed as explained above and the POI was eluted in a linear gradient to 400 mM imidazole. Pure fractions were pooled and concentrated and applied to HiLoad 16/600 Superdex 200 pg (Cytiva) in a buffer with 2.5% glycerol.

Generation of the CK2 holoenzyme was loosely based on previous literature (*104*, *105*). The CK2α^1-335^ expression plasmid carried a resistance against Kanamycin, while the CK2β^1-193^ expression plasmid was resistant against Ampicillin. The expression was performed in *E. coli* BL21(DE3)-Codon-plus-RIL in TB medium with the specific antibiotic and additional Chloramphenicol. The cells were grown to an OD_600_ of 0.6 at 37 °C. The expressions were induced by the addition of 0.5 mM IPTG and incubated at 30 °C for 4 h. The two isolated cultures were harvested and mixed together. The cells were resuspended in lysis buffer (50 mM Tris pH 8.5, 500 mM NaCl, 30 mM imidazole). The centrifuged lysate was incubated at 4 °C for 16 h to ensure the efficient formation of the holoenzyme. The next day, the lysate was filtered and purified on a 5 ml HisTrap FF (Cytiva). The column was washed after the application of the lysate and protein was eluted in lysis buffer with 250 mM imidazole. The eluate was concentrated and applied to SEC on a HiLoad 16/600 Superdex 200 pg (Cytiva) in SEC buffer (25 mM Tris pH 8.5, 500 mM NaCl).

### Analytical SEC

Proteins were mixed in SEC buffer (20 mM HEPES pH 6.8, 300 mM NaCl, 2.5% glycerol, 1 mM TCEP), diluted to 5 µM in 55 µl. Complexes were incubated for at least 1 h at 4 °C. Samples were centrifuged and 5 µl of sample were taken for SDS-PAGE analysis prior to SEC on a Superose 6 Increase 5/150 GL (Cytiva, Marlborough, US-MA) on an ÄKTA micro system (Cytiva). All samples were eluted under isocratic conditions at 4 °C in SEC buffer at a flow rate of 0.2 ml/min. Fractions of 100 µl were collected and analyzed by SDS-PAGE and Coomassie blue staining.

### Pull-down assays

The proteins were mixed at 3 µM in binding buffer (20 mM HEPES pH 6.8, 300 mM NaCl, 2.5% glycerol, 1 mM TCEP, 0.01% Tween) to a total volume of 50 µl. The samples were incubated at 4 °C for at least 1 h. Afterwards, they were centrifuged for 15 min at 16,000 xg prior to mixing them with 25 µl of amylose beads (New England Biolabs) or glutathione beads (Serva). Then, 20 µl were taken as an input sample for SDS-PAGE. The rest of the solution was incubated at 4 °C for an additional hour on an orbital shaker set to 1000 rpm (IKA VXR basic Vibrax). The samples were centrifuged, at 800 xg for 3 min at 4 °C. The unbound protein in the supernatant was removed, and the beads were washed 4 times with 500 µl of binding buffer. At the last step, the maximum amount of buffer was carefully removed and beads were taken up in 20 µl of SDS-PAGE sample loading buffer. The samples were boiled for 5 min at 96 °C and analyzed by SDS-PAGE and Coomassie staining.

### *In vitro* phosphorylation

Proteins were diluted to 2.5 µM in 20 mM Hepes pH 6.8, 300 mM NaCl, 2.5% glycerol and 1 mM TCEP. Otherwise, concentrations and buffer were used according to analytical SEC or pull-down assays. 1 mM Sodium orthovanadate and 5 µM okadaic acid were added to inhibit residual Lambda-phosphatase activity and 10 mM MgCl_2_ and 2 mM ATP were added for kinase activity. The samples were incubated at 25 °C for 90 minutes with CK2 at a 1:20 ratio. Pro-Q Diamond phosphoprotein stain (Invitrogen, Carlsbad, California, United States) was performed according to the manufacturer’s manual.

### Identification of phosphorylation sites by mass spectrometry

Liquid chromatography coupled to mass spectrometry was used to analyze phosphorylation sites on FACT, phosphorylated *in vitro* by CK2 as described above. Insect cell-expressed and dephosphorylated FACT were subjected to the analysis as controls. Samples were reduced, alkylated and digested with LysC/Trypsin and prepared for mass spectrometry as previously described (*106*).

### Cell culture

All Cells were grown at 37 °C in the presence of 5% CO_2_. Parental Flp-In T-Rex DLD-1 osTIR1 and DLD-1^YFP-AID^ cells were a kind gift from D. C. Cleveland (University of California, San Diego, USA). DLD-1 cells were grown in Dulbecco’s modified Eagle’s medium (DMEM; PAN Biotech) and parental hTERT RPE-1 cells and hTERT RPE-1 cells expressing endogenously tagged CENP-U-FKBP^F36V^ were grown in DMEM F12 media (PAN Biotech); both supplemented with 10% tetracycline-free fetal bovine serum (Sigma) and l-glutamine (PAN Biotech).

### Cell synchronisation and drug treatments

Degradation of the endogenous CENP-C^YFP-AID^ was achieved through the addition of 500 μM Indole-Acetic-Acid (IAA, Sigma-Aldrich) and degradation of endogenous CENP-U-FKBP^F36V^was achieved by addition of 500 nM dTAG^V^-1 (Tocris). Cells were synchronized using S-trityl-L-cysteine (STLC) at 5 µM for 16 hours or Nocodazole 3.3 µM for 4 hours.

### RNA interference

Depletion of endogenous proteins was achieved through transfection of small interfering RNA (siRNA) with RNAiMAX (Invitrogen) according to manufacturer’s instructions. Following siRNAs treatments were performed in this study: 30 nM of each oligo, siCENP-T (Dharmacon, 5′-GACGAUAGCCAGAGGGCGU-3′, 5’-AAGUAGAGCCCUUACACGA-3’) for 60 hours (*74*), siCENP-H (Sigma, 5′-CUAGUGUGCUCAUGGAUAA-3′) (*64*), siCENP-I (Sigma, 5′-AAGCAACTCGAAGAACATCTC-3′) (*107*), siCENP-K (Dharmacon, On-target plus smartpool-XX) (*73*), siCENP-M (Sigma, 5′-ACAAAAGGUCUGUGGCUAA-3′, 5’-UUAAGCAGCUGGCGUGUUA-3’, 5’-GUGCUGACUCCAUAAACAU-3’) (*74*) for 72 hours,

### Generation of stable cell lines

Stable Flp-In T-Rex DLD-1 osTIR1 cell lines were generated using FRT/Flp recombination. CENP-U^fl^ and 115-C constructs were cloned into pcDNA5 plasmids (*75*) and were co-transfected with pOG44 (Invitrogen), encoding the Flp recombinase, into DLD-1 cells using X-tremeGENE (Roche) according to the manufacturer’s instructions. After selection for 2 weeks in DMEM supplemented with hygromycin B (250 μg/ml; Carl Roth) and blasticidin (4 μg/ml; Thermo Fisher Scientific), single-cell colonies were isolated, expanded and the expression of the transgenes was checked by immunofluorescence microscopy and immunoblotting analysis. The gene expression was induced by the addition of 0.3 μg/ml doxycycline (Sigma-Aldrich).

hTERT RPE-1 CENP-U-FKBP^F36V^-NeoR knock-in cell line was generated via electroporation of gRNA-Cas9 ribonucleoproteins (RNPs) as previously described (*108*). Briefly, 3×10^5^ cells were electroporated using P3 Primary Cell Nucleofector^®^ 4D Kit and Nucleofector 4D system (Lonza) with 400 ng donor DNA, 120 pmol of Cas9, 1.5 µl Alt-R™-CRISPR-Cas9 crRNA (TTAGAGAAGCTCCTTGACCA (*GGG*)), 1.5 μl Alt-R^®^ CRISPR-Cas9 tracrRNA (100 μM, IDT) and 1.2 μl of Alt-R^®^ Cas9 Electroporation enhancer (100 μM, IDT). After electroporation, cells were seeded into DMEM F12 media supplemented with 1 µM NU7441 for 48 h before selection with 400 µg/ml of G418 for 2 weeks. The pool of cells was subjected to single cell dilution to obtain monoclonal lines. Genomic DNA was isolated from clones and in-frame knock-in was confirmed by Sanger sequencing using primers spanning the locus of insertion (Primer fwd: 5’-CATGTGTGTGGTAGTCACAGCATG-3’, Primer rev: 5’-TCTGGGATAATGGCATTGATGATGC-3’).

### Immunofluorescence

Cells were grown on coverslips pre-coated with poly-L-lysine (Sigma-Aldrich). Cells were pre-permeabilized with 0.5% Triton X-100 solution in PHEM (Pipes, HEPES, EGTA, MgCl_2_) buffer supplemented with 100 nM microcystin for 5 minutes before fixation with 4% paraformaldehyde (PFA) in PHEM for 15 minutes. After blocking with 5% boiled goat serum (BGS) in PHEM buffer for 30 min, cells were incubated for 2 hours at room temperature with the following primary antibodies; CENP-C (Guinea pig, MBL, #PD030, 1:1,000), CENP-HK, CENP-TW (Rabbit, made in-house, 1:800), SSRP1 (Mouse, BioLegend Europe #609702, 1:200), CENP-A (Mouse, GeneTex GTX13939, 1:500), CENP-O (Gift from McAinsh Lab, 1:200), PLK1 (Mouse,Abcam #ab17057, 1:500), CENP-R (Rabbit, Proteintech #107431-AP, 1:200), CREST (Anticentromere Anti immune serum) (Human, Antibodies Inc. (via antibodies-online), #15-234, 1:200) diluted in 2.5% BGS-PHEM with exception of CENP-U (Sigma (Atlas, HPA022048, 1:100) which was diluted in 5% BGS-PHEM and incubated at 37 °C for 3 hours.

Subsequently, cells were incubated for 1 hour at room temperature with the following secondary antibodies: (all 1:200 in 2.5% BGS-PHEM): Goat anti-mouse Alexa Fluor 488 (Invitrogen A A11001), goat anti-mouse Rhodamine Red (Jackson Immuno Research 115-295-003), donkey anti-rabbit Alexa Fluor 488 (Invitrogen A21206), donkey anti-rabbit Rhodamine Red (Jackson Immuno Research 711-295-152), goat anti-human Alexa Fluor 647 (Jackson Immuno Research 109-603-003), goat anti-guinea pig Alexa Fluor 647 (Invitrogen A-21450). All washing steps were performed with PHEM supplemented with 0.1% Triton-X-100 (PHEM-T) buffer. DNA was stained with 0.5 μg/ml DAPI (Serva) and Mowiol (Calbiochem) was used as mounting media.

Cells were imaged at room temperature using a spinning disk confocal device on the 3i Marianas system equipped with an Axio Observer Z1 microscope (Zeiss, Jena, Germany), a CSU-X1 confocal scanner unit (Yokogawa Electric Corporation, Tokyo, Japan), 100x / 1.4NA oil objectives (Zeiss), and Orca Flash 4.0 V2 sCMOS Camera (Hamamatsu). Images were acquired as z sections at 0.27 μm using Slidebook Software 2023.3 (Intelligent Imaging Innovations, Denver, USA). Images were converted into maximum intensity projections and exported as 16-bit TIFF files. Alternatively, cells were imaged using a UPLSAPO 100x / 1.4NA oil objective on a DeltaVision deconvolution microscope (GE Healthcare, UK) equipped with an IX71 inverted microscope (Olympus, Japan), and a pco.edge sCMOS camera (PCO-TECH Inc., USA). Images were acquired as z sections at 0.2 μm

Quantification of kinetochore signals were performed on 16-bit maximum intensity projections using a semi-automatic quantification MACRO in FIJI (*109*) with background subtraction. Data was exported to Microsoft Excel for normalization and plotted using GraphPad Prism 9 software. Statistical analysis was performed with a non-parametric t-test comparing two unpaired groups (Mann-Whitney test). Symbols indicate n.s. p>0.05, * p:<0.05, ** p:<0.01, ***p:<0.001, ****p:<0.0001

Images were assembled in Adobe Illustrator 2024.

### Co-immunoprecipitation

For co-immunoprecipitaion, DLD-1 cells expressing EGFP, ^EGFP^CENP-U^fl^ and ^EGFP^CENP-U^115-C^ were harvested by mitotic shake off and lysed using lysis buffer (75 mM HEPES pH 7.5, 150 mM KCl, 10% glycerol, 1 mM EGTA, 1.5 mM MgCl_2_, 1 mM DTT, 0.075% NP-40) supplemented with 1 mM PMSF, Protease-Inhibitor Mix HP Plus (Serva), phosphatase inhibitor PhosSTOP (Sigma Aldrich), and Benzoase (EMD Millipore Corp,USA)). Lysates were clarified by centrifugation at 22,000 g for 30 min at 4 °C and supernatant was collected for immunoprecipitation analysis. Lysates at a concentration of 7 mg/ml were incubated with 20 µl GFP-Trap magnetic agarose (ChromoTEK) for 3 hours at 4 °C in a total volume of 500 µl in Lysis Buffer. Beads were washed 3 times with Lysis Buffer. The dry beads were resuspended in SDS-PAGE sample loading buffer and boiled for 5 minutes at 95 °C. The samples were analyzed by SDS-PAGE and subsequent Western blotting analysis. The following antibodies were used: GFP (rabbit, made in-house, 1:3,000), SSRP1 (Mouse, BioLegend, 1:1000) PLK1 (1:1000.), anti-mouse or anti-rabbit (1:10,000; Amersham, NXA931 and NA934) conjugated to horseradish peroxidase were used. After incubation with ECL Western blotting reagent (GE Healthcare), images were acquired with the ChemiDoc MP System (Bio-Rad) using Image Lab 6.0.1 software.

## Acknowledgments

We thank Franziska Müller and Petra Janning for help with mass spectrometry experiments, Dongqing Pan for the generation of initial constructs for the expression of recombinant FACT, Karsten Niefind for providing expression constructs for CK2, Duccio Conti for the help in initial experiments, Sabine Wohlgemuth for the purification of CENP-LN, Carolin Körner for the preparation of recombinant kinases, Lia Nitz for the production of a subset of ^MBP^CENP-C fusion proteins, Nico Schmidt for help with microscopy experiments and data analysis, and Daniele Fachinetti and Don C. Cleveland for sharing cell lines.

## Author contributions

Conceptualization: JS, BM, AM; Investigation: JS, BM, IH, DV, MEP; Visualization: JS, BM, AM; Supervision: AM; Writing—original draft: JS, AM; Writing—review & editing: all authors.

## Data and materials availability

All data needed to evaluate the conclusions in the paper are present in the paper and the Supplementary Materials.

## Supplementary Materials

**Figure S1.**
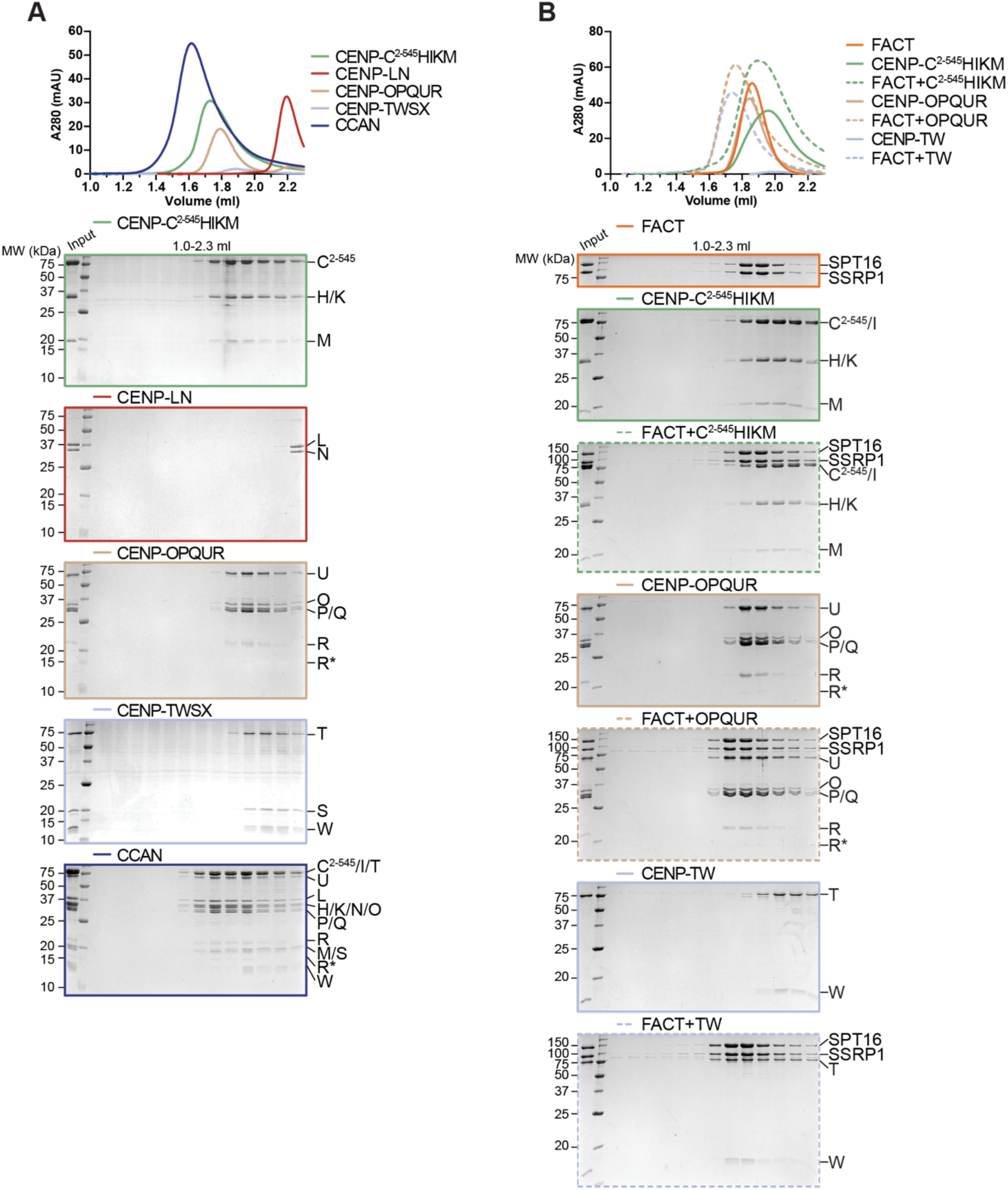
(related to Figure 1) **(A)** Analytical SEC of the individual CCAN subcomplexes and its reconstitution. **(B)** Analytical SEC of FACT and CENP-C^2-545^HIKM, CENP-OPQUR and CENP-TW with Coomassie-stained SDS-PAGEs below.

**Figure S2.**
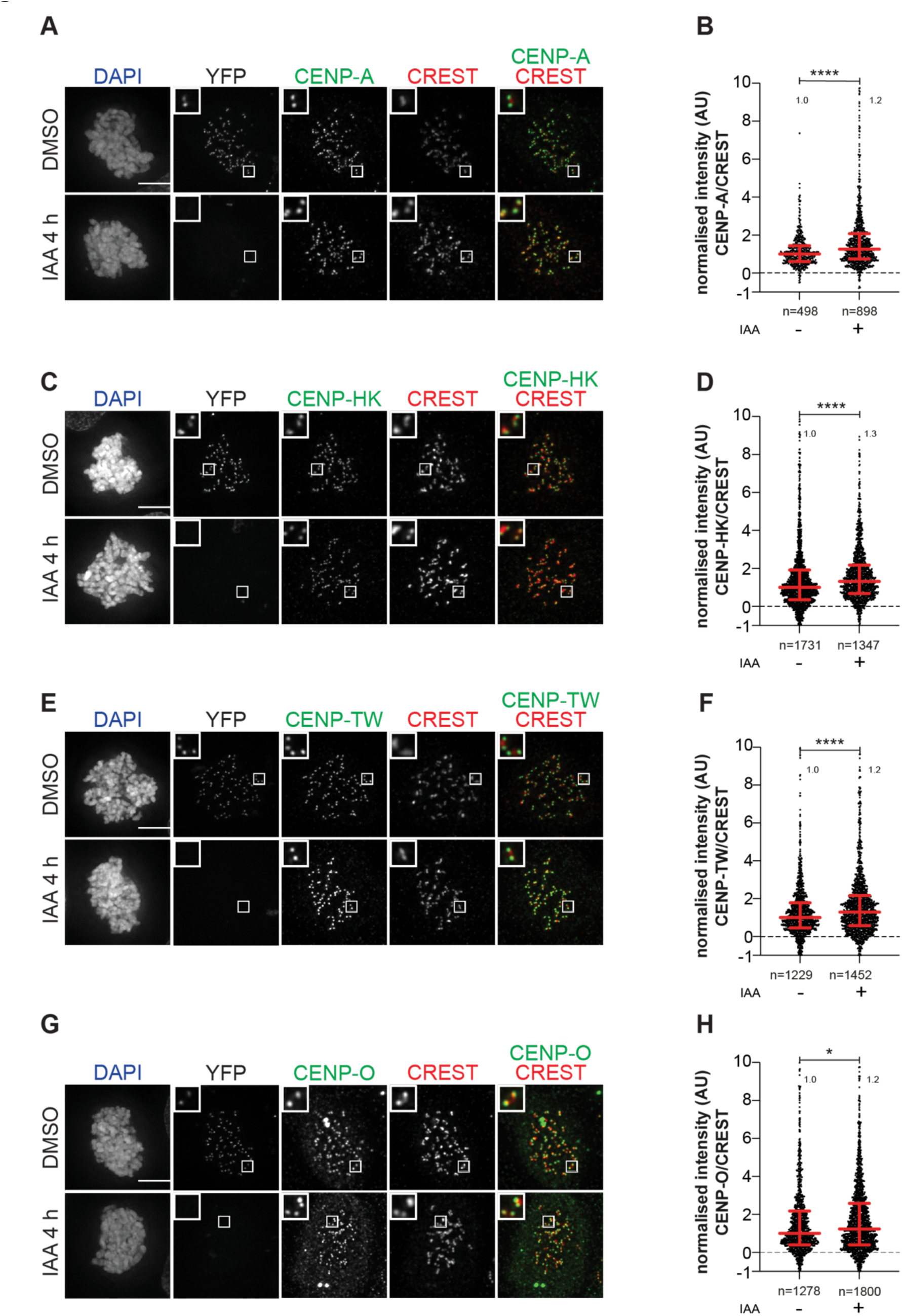
(related to Figure 1) **(A)** Representative images of localization of CENP-A after degradation of CENP-C in DLD-1-CENP-C^YFP-AID^ cells for 4 hours. Cells were treated with IAA (500 µM) to degrade endogenous CENP-C and Nocodazole (3.3 µM) to get mitotic population of cells for 4 hours. CREST serum was used to visualize kinetochores and DAPI to stain DNA. Three biological replicates were performed. Scale bar: 5 μm. **(B)** Scatter plot of CENP-A levels at kinetochores for the experiment shown in panel (A). *n* is the number of individually measured kinetochores. **(C)** Representative images of localization of CENP-HK after degradation of CENP-C in DLD-1-CENP-C^YFP-AID^ cells for 4 h. Cells were treated with IAA (500 µM) to degrade endogenous CENP-C and Nocodazole (3.3 µM) to get mitotic population of cells for 4 hours. CREST serum was used to visualize kinetochores and DAPI to stain DNA. Three biological replicates were performed. Scale bar: 5 μm. **(D)** Scatter plot of CENP-HK levels at kinetochores for the experiment shown in panel (C). *n* refers to individually measured kinetochores. **(E)** Representative images of localization of CENP-TW after degradation of CENP-C in DLD-1-CENP-C^YFP-AID^ cells for 4 hours. Cells were treated with IAA (500 µM) to degrade endogenous CENP-C and with Nocodazole (3.3 µM) for 4 hours to enrich for mitotic cells. CREST serum was used to visualize kinetochores and DAPI to stain DNA. Three biological replicates were performed. Scale bar: 5 μm. **(F)** Scatter plot of CENP-TW levels at kinetochores for the experiment shown in panel (E). *n* refers to individually measured kinetochores. **(G)** Representative images of localization of CENP-O after degradation of CENP-C in DLD-1-CENP-C^YFP-AID^ cells for 4 hours. Cells were treated with IAA (500 µM) to degrade endogenous CENP-C and Nocodazole (3.3 µM) to get mitotic population of cells for 4 hours. CREST was used to visualize kinetochores and DAPI to stain DNA. Three biological replicates were performed. Scale bar: 5 μm. **(H)** Scatter plot of CENP-O levels at kinetochores for the experiment in panel (G). *n* is the number of individually measured kinetochores.

**Figure S3.**
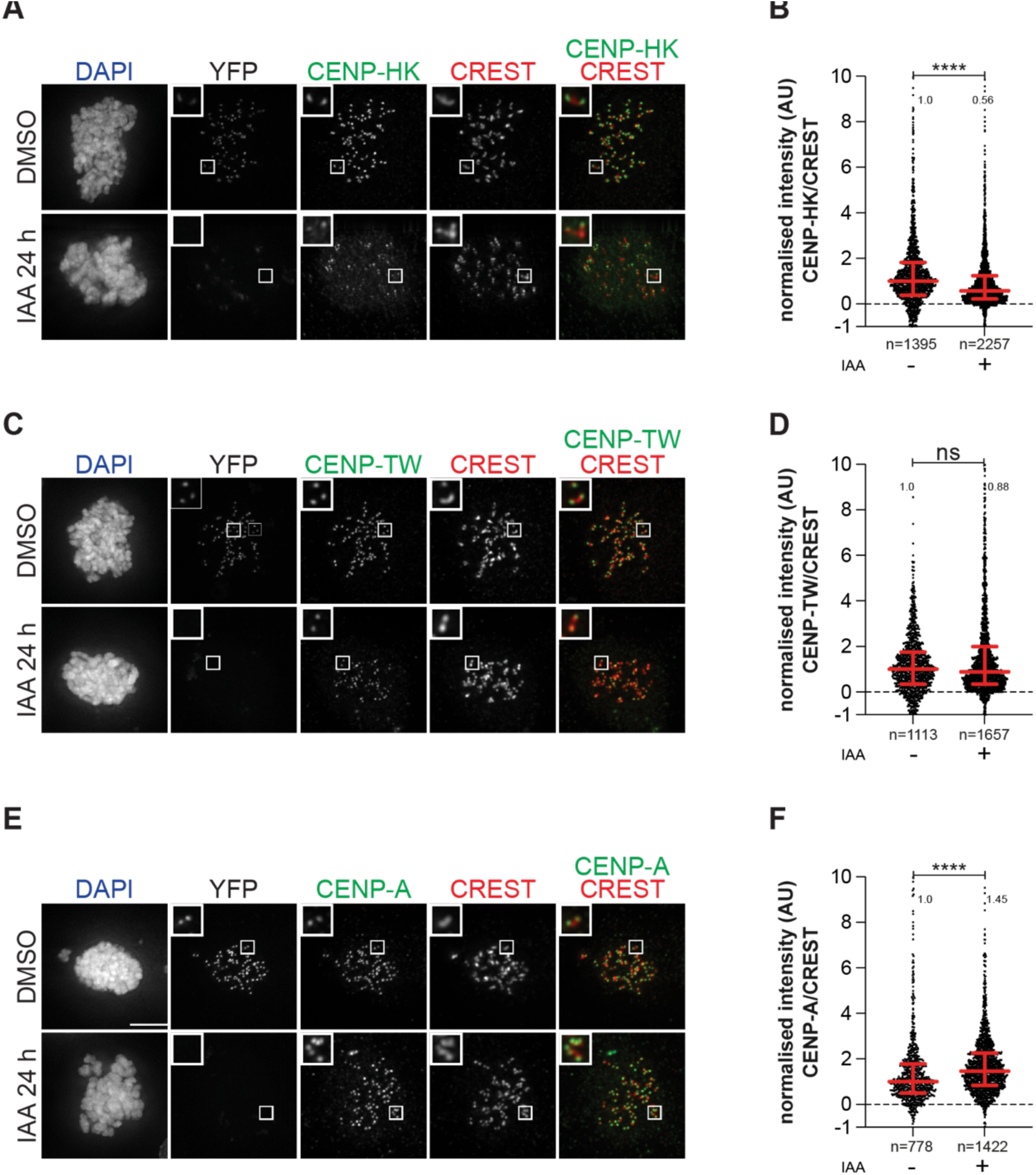
(related to Figure 1) **(A)** Representative images of localization of CENP-HK after degradation of CENP-C in DLD-1-CENP-C^YFP-AID^ cells for 24 hours. Cells were treated with IAA (500 µM) to degrade endogenous CENP-C for 24 hours and Nocodazole (3.3 µM) for 4 hours to get mitotic population of cells. CREST was used to visualize kinetochores and DAPI to stain DNA. Three biological replicates were performed. Scale bar: 5 μm. **(B)** Scatter plot of CENP-HK levels at kinetochores of the experiment shown in panel (A). *n* is the number of individually measured kinetochores. **(C)** Representative images of localization of CENP-TW after degradation of CENP-C in DLD-1-CENP-C^YFP-AID^ cells for 24 hours. Cells were treated with IAA (500 µM) to degrade endogenous CENP-C for 24 hours and Nocodazole (3.3 µM) for 4 hours to get mitotic population of cells. CREST serum was used to visualize kinetochores and DAPI to stain DNA. Three biological replicates were performed. Scale bar: 5 μm. **(D)** Scatter plot of CENP-TW levels at kinetochores of the experiment shown in panel (C). *n* is the number of individually measured kinetochores. **(E)** Representative images of localization of CENP-A after degradation of CENP-C in DLD-1-CENP-C^YFP-AID^ cells for 24 hours. Cells were treated with IAA (500 µM) to degrade endogenous CENP-C for 24 hours. Nocodazole (3.3 µM) was added for 4 hours to enrich for mitotic cells. CREST serum was used to visualize kinetochores and DAPI to stain DNA. Three biological replicates were performed. Scale bar: 5 μm. **(F)** Scatter plot of CENP-A levels at kinetochores of the experiment shown in panel (E). *n* is the number of individually measured kinetochores.

**Figure S4.**
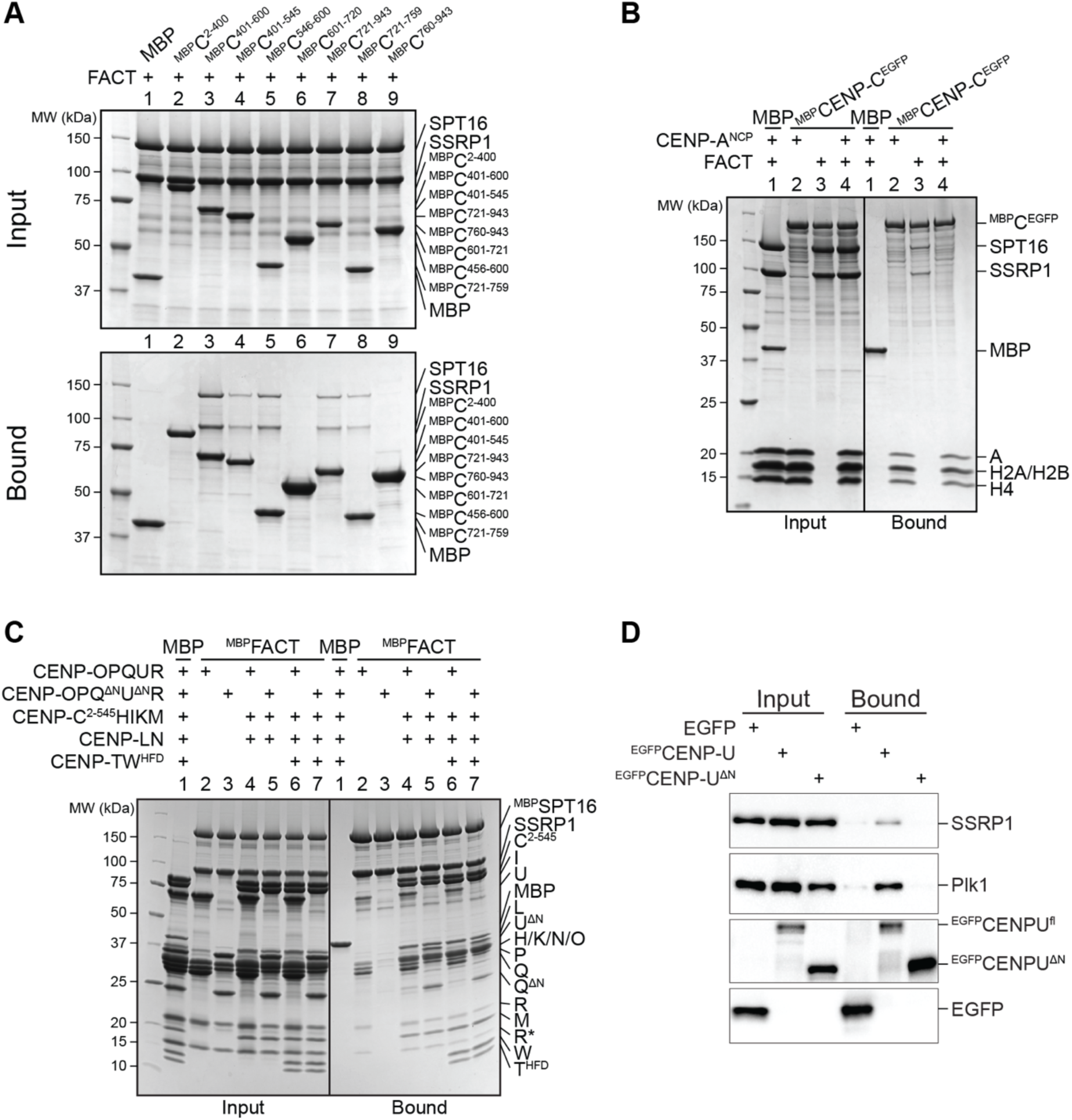
(related to Figure 2) **(A)** Amylose-resin pull-down assay with a set of ^MBP^CENP-C fusion proteins as baits spanning the entire sequence of CENP-C and FACT as a prey. **(B)** Amylose-resin pull-down assay using immobilized ^MBP^CENP-C^EGFP^ on beads and adding FACT and CENP-A^NCP^ as preys. CENP-A^NCP^ is the histone octamer reconstituted on 145-bp Widom 601 sequence. **(C)** Amylose-resin pull-down assay with ^MBP^FACT as a bait to analyze the influence of the CENP-QU N-terminal tails on the binding of CENP-OPQUR, -C^2-545^HIKMLNOPQUR and -C^2-545^HIKMLNOPQURTW^HFD^. **(D)** Lysates prepared from STLC synchronized DLD-1 cells expressing GFP alone or GFP-CENP-U Fl or 115-C were subjected to immunoprecipitation using GFP-trap beads followed by Western blotting with antibodies against GFP, SSRP1 and PLK1. PLK1 was used as internal control.

**Figure S5.**
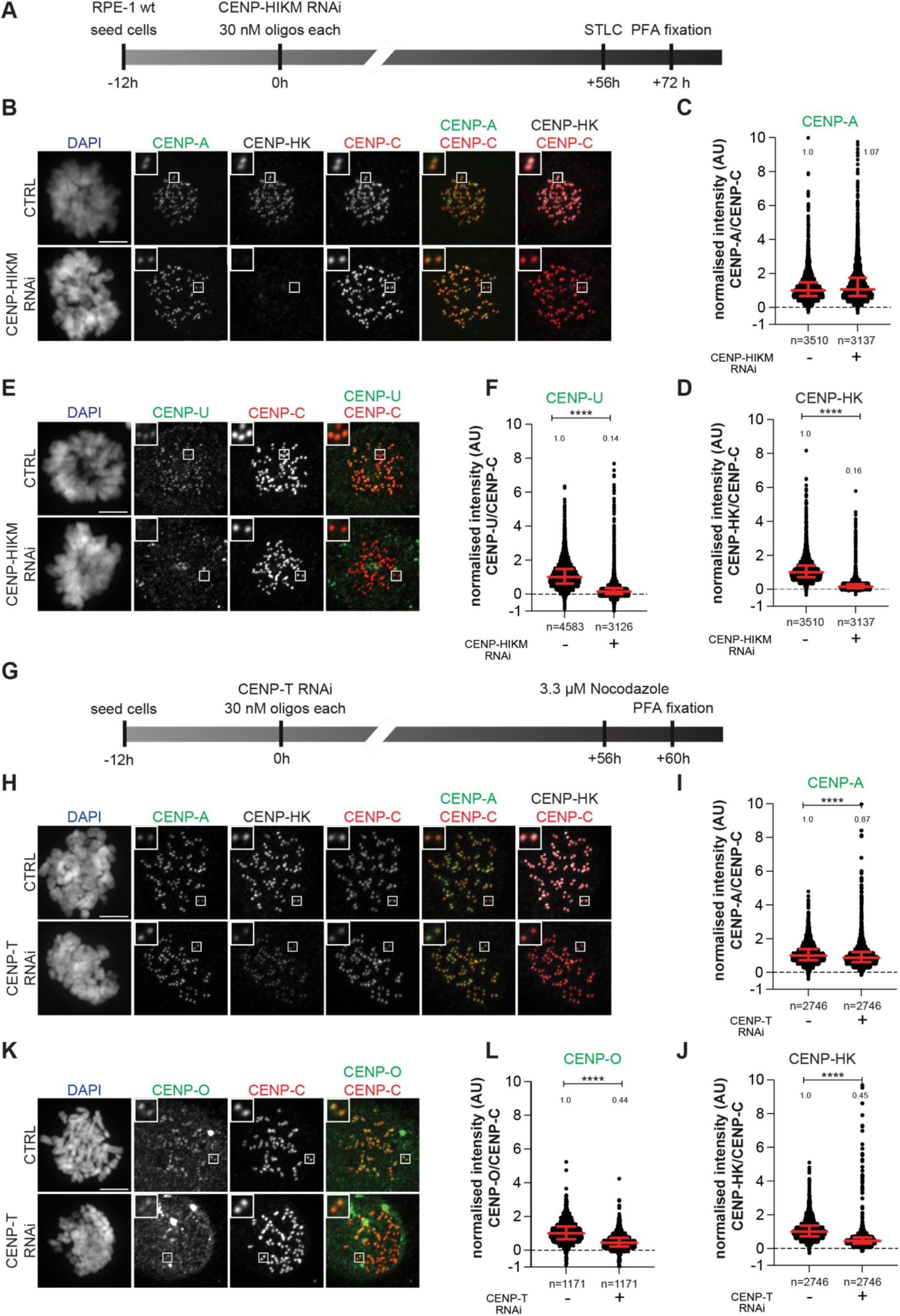
(related to Figure 3) **(A)** Schematic representation of experimental scheme used for CENP-HIKM RNAi **(B)** Representative images of localization of CENP-A and CENP-HK after depletion of CENP-HIKM complex in RPE-1 cells. CENP-HIKM RNAi was performed for 72 hours using oligos for each subunit at 30 nM concentration. Cells were treated with STLC (5 µM) for 16 hours to obtain a mitotic population before fixation. CENP-C was used to visualize kinetochores and DAPI to stain DNA. Three biological replicates were performed. Scale bar: 5 μm. **(C)** Scatter plot of CENP-A levels at kinetochores for the experiment shown in panel (B). *n* is the number of individually measured kinetochores. **(D)** Scatter plot of CENP-HK levels at kinetochores for the experiment in panel (B). *n* is the number of individually measured kinetochores. **(E)** Representative images of localization of CENP-U after depletion of CENP-HIKM complex in RPE-1 cells. CENP-HIKM RNAi was performed for 72 hours using oligos for each subunit at 30 nM concentration. To obtain mitotic cells, cells were treated with STLC (5 µM) for 16 hours before fixation. CENP-C visualizes kinetochores and DAPI stains DNA. Three biological replicates were performed. Scale bar: 5 μm. **(F)** Scatter plot of CENP-U levels at kinetochores of the experiment shown in panel (E). *n* refers to individually measured kinetochores. **(G)** Schematic representation of experimental scheme used for CENP-T RNAi **(H)** Representative images of localization of CENP-A and CENP-HK after depletion of CENP-T complex in RPE-1 cells. CENP-T RNAi was performed for 60 hours using oligos for each subunit at 30 nM concentration. Cells were treated with Nocodazole (3.3 µM) for 4 hours before fixation to enrich for mitotic cells. CENP-C visualizes kinetochores and DAPI stains DNA. Three biological replicates were performed. Scale bar: 5 μm. **(I)** Scatter plot of CENP-A levels at kinetochores for the experiment in panel (H). *n* is the number of individually measured kinetochores. **(J)** Scatter plot of CENP-HK levels at kinetochores for the experiment in panel (H). *n* is the number of individually measured kinetochores. **(K)** Representative images of localization of CENP-O after depletion of CENP-T complex in RPE-1 cells. CENP-T RNAi was performed for 60 hours using oligos for each subunit at 30 nM concentration. Cells were treated with Nocodazole (3.3 µM) for 4 hours before fixation to obtain mitotic population. CENP-C visualizes kinetochores and DAPI stains DNA. Three biological replicates were performed. Scale bar: 5 μm. **(L)** Scatter plot of CENP-O levels at kinetochores for the experiment in panel (K). *n* is the number of individually measured kinetochores.

**Figure S6.**
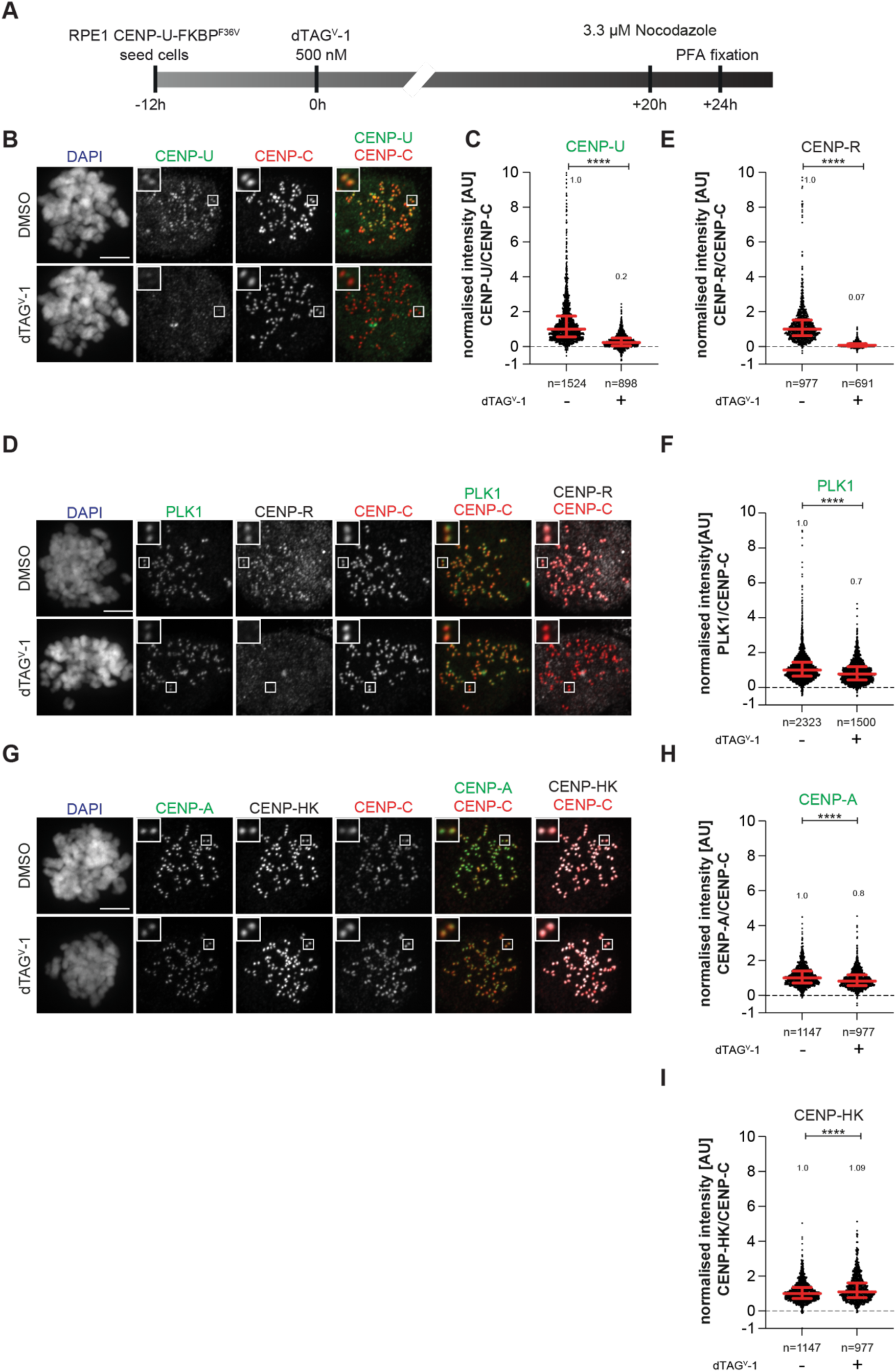
(related to Figure 3) **(A)** Schematic representation of experimental scheme used for CENP-U dTAG^V^-1 treatment **(B)** Representative images of localization of CENP-U after depletion of CENP-U complex in RPE-1-CENP-U-FKBP^F36V^ cells. Cells were treated with dTAG^V^-1 (500 nM) for 24 hours to degrade endogenous CENP-U complex. Cells were treated with Nocodazole (3.3 µM) for 4 hours to obtain a mitotic cell population prior to fixation. CENP-C was used to visualize kinetochores and DAPI to stain DNA. Three biological replicates were performed. Scale bar: 5 μm. **(C)** Scatter plot of CENP-U levels at kinetochores for the experiment in panel (B). *n* is the number of individually measured kinetochores. **(D)** Representative images of localization of CENP-R and PLK1 after depletion of CENP-U complex in RPE-1-CENP-U-FKBP^F36V^ cells. Cells were treated with dTAG^V^-1 (500 nM) for 24 hours to degrade endogenous CENP-U complex. Cells were treated with Nocodazole (3.3 µM) for 4 h prior to fixation to obtain mitotic population. CENP-C was used to visualize kinetochores and DAPI to stain DNA. Three biological replicates were performed. Scale bar: 5 μm. **(E)** Scatter plot of CENP-R levels at kinetochores for the experiment in panel (D). *n* is the number of individually measured kinetochores. **(F)** Scatter plot of PLK1 levels at kinetochores for the experiment in panel (D). *n* is the number of individually measured kinetochores. **(G)** Representative images of localization of CENP-A and CENP-HK after depletion of CENP-U complex in RPE-1-CENP-U-FKBP^F36V^ cells. Cells were treated with dTAG^V^-1 (500 nM) for 24 hours to degrade endogenous CENP-U complex. Cells were treated with Nocodazole (3.3 µM) for 4 h prior to fixation to obtain mitotic population. CENP-C was used to visualize kinetochores and DAPI to stain DNA. Three biological replicates were performed. Scale bar: 5 μm. **(H)** Scatter plot of CENP-A levels at kinetochores of the experiment shown in panel (G). *n* is the number of individually measured kinetochores. **(I)** Scatter plot of CENP-HK levels at kinetochores of the experiment shown in panel (G). *n* is the number of individually measured kinetochores.

**Figure S7.**
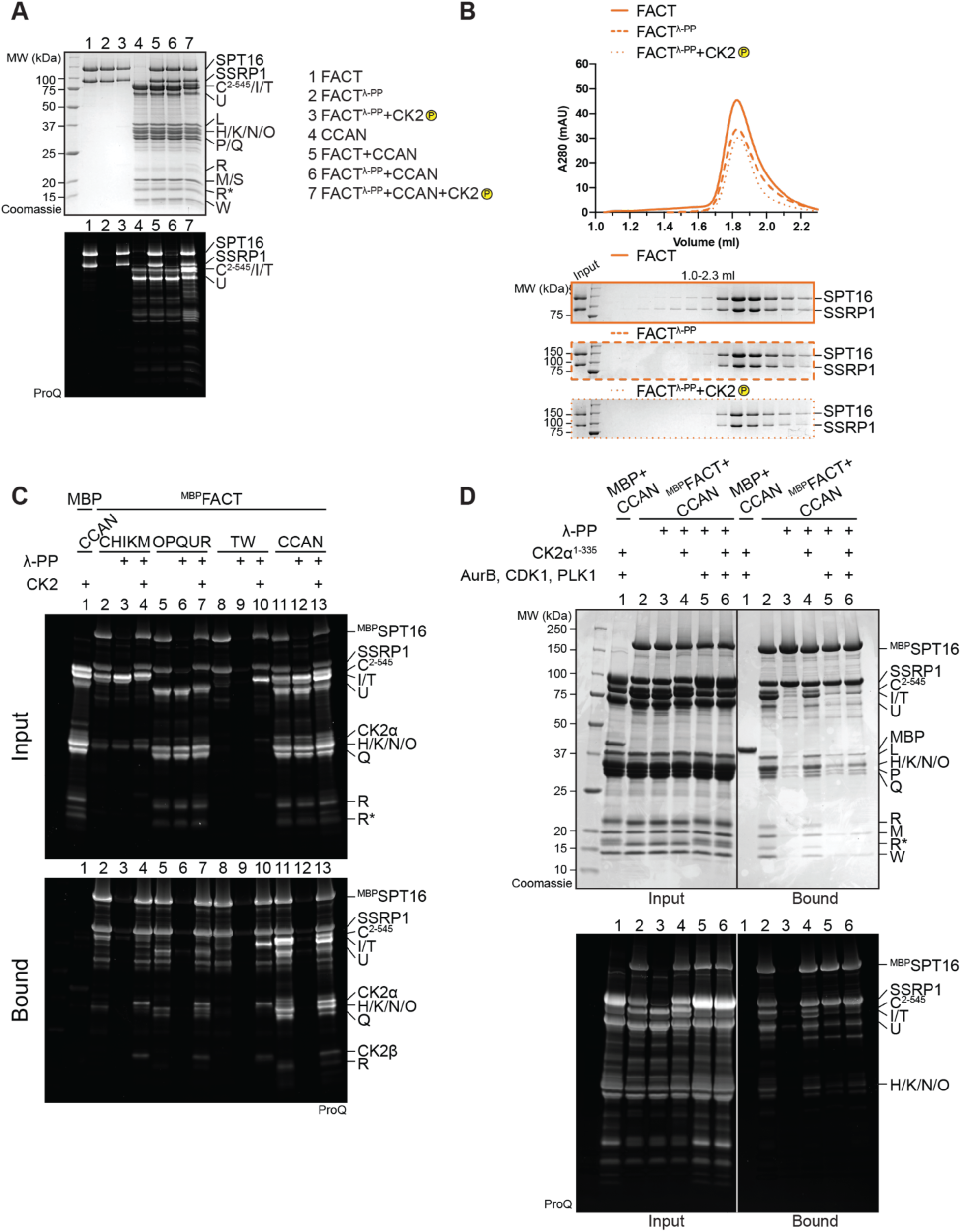
(related to Figure 5) **(A)** ProQ Diamond staining to monitor the phosphorylation state of samples in Figure 5A. **(B)** Additional samples of the analytical SEC experiment in Figure 5A verify that the phosphorylation state of FACT does not alter its elution volume. **(C)** ProQ Diamond staining of the pull-down assay in Figure 5B. **(D)** Amylose-resin pull-down assay to analyze CCAN binding to dephosphorylated ^MBP^FACT upon phosphorylation by CK2α^1-335^ or Aurora B, CDK1 and PLK1 or all of them. CDK1 indicates the use of a complex of CDK1/Cyclin-B/CKS1. The ProQ Diamond staining of the SDS-PAGE is shown below.

**Figure S8.**
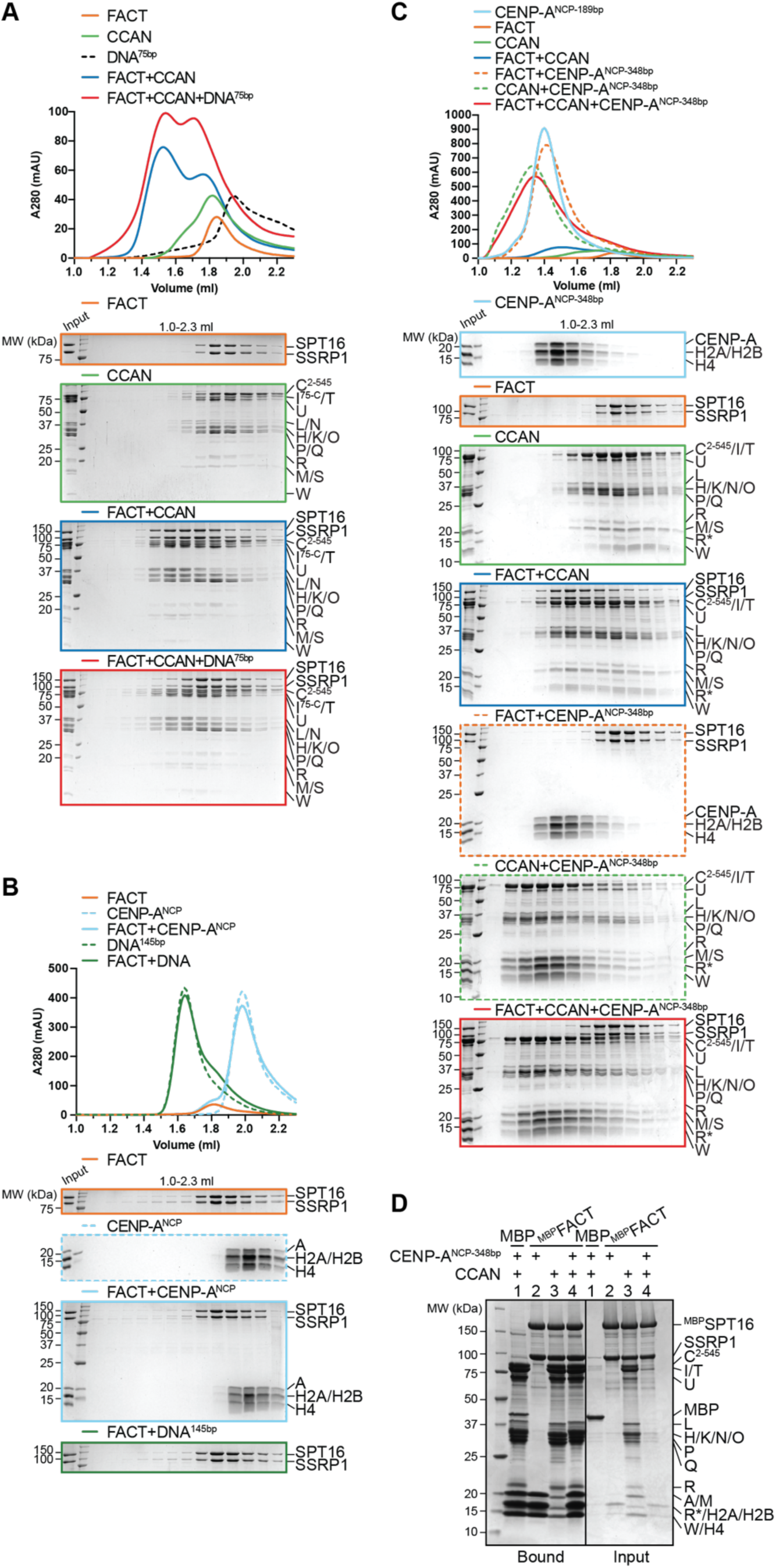
(related to Figure 6) **(A)** Analytical SEC of FACT and CCAN upon the addition of a 75-bp CEN1-like DNA. **(B)** Analytical SEC to test binding of FACT to CENP-A^NCP^ or 145-bp Widom 601 DNA. **(C)** Analytical SEC of FACT, CCAN, and CENP-A^NCP^ on a 348-bp DNA. The histone octamer was reconstituted on a 183-bp CEN1-like sequence and 165 CEN1-like sequence was ligated to it. **(D)** Amylose-resin pull-down assay of ^MBP^FACT and CCAN upon addition of CENP-A^NCP-348bp^.

**Figure S9.**
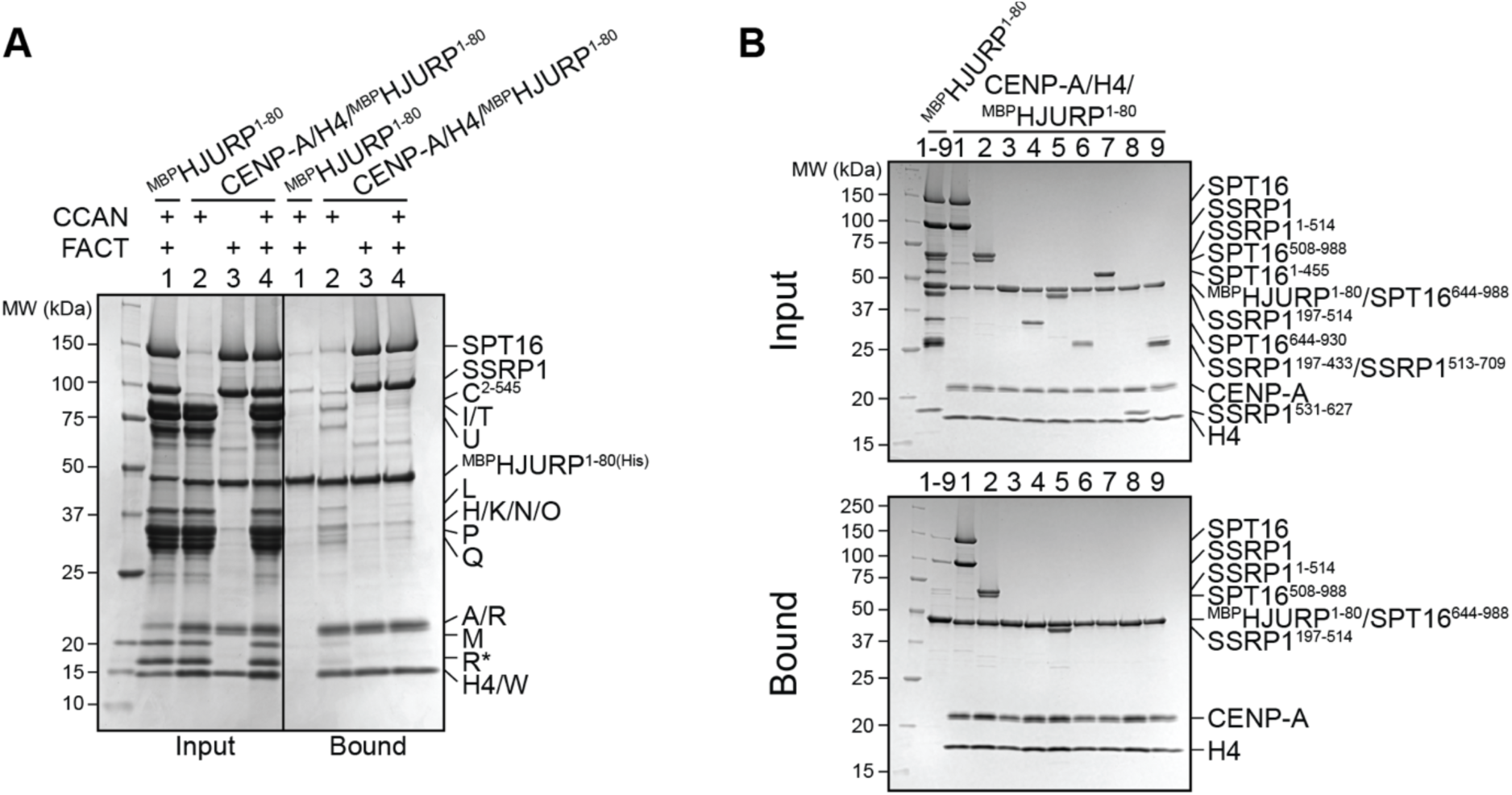
(related to Figure 6) **(A)** Amylose-resin pull-down assay using ^MBP^HJURP^1-80^ in complex with CENP-A/H4 as bait and CCAN and FACT as preys. ^MBP^HJURP^1-80^ in absence of histones was used as a negative control. **(B)** Amylose-resin pull-down assay using ^MBP^HJURP^1-80^ in complex with CENP-A/H4 as bait and FACT constructs (Fig. 4A) as preys. ^MBP^HJURP^1-80^ in absence of histones was used as a negative control.

**Table S1.**
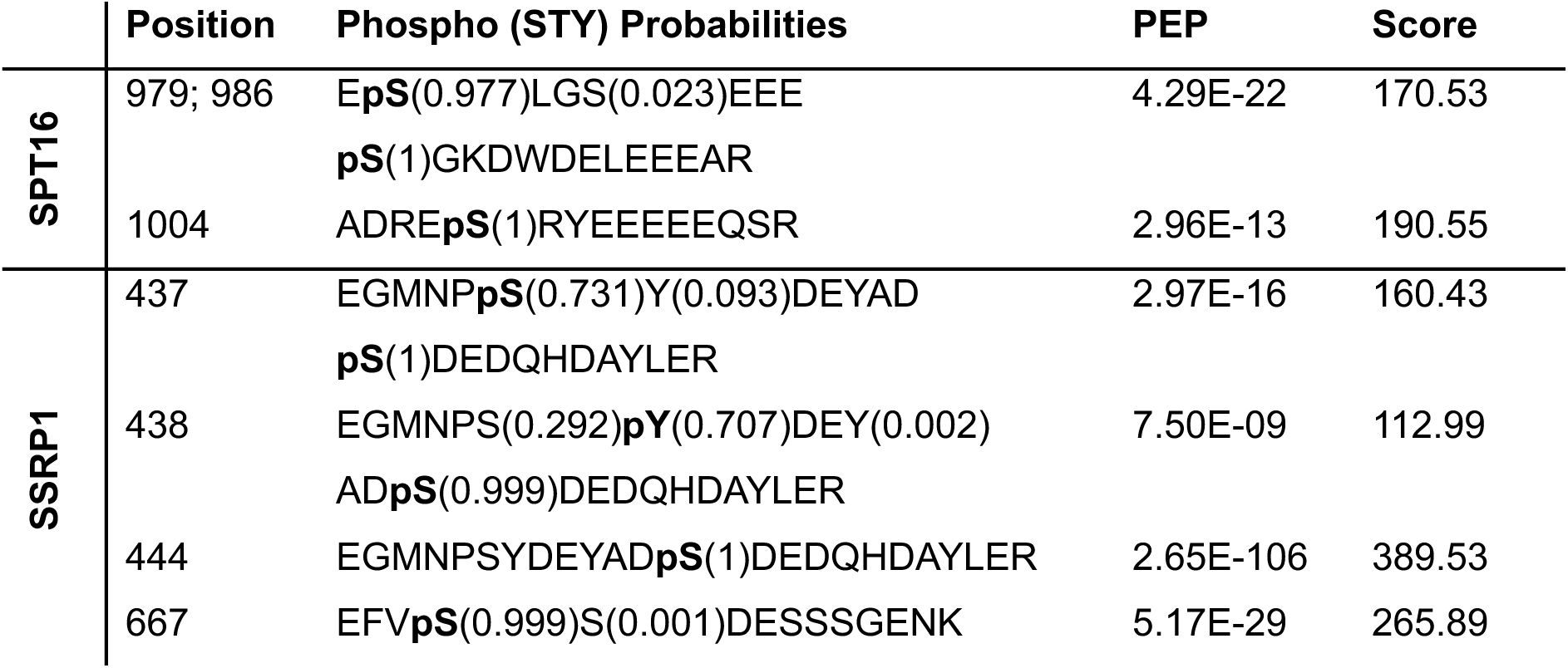
Phospho-sites on FACT detected by mass spectrometry subsequent to treatment with CK2. Sites in bold followed by the probability, the p-value of the posterior error probability (PEP) of the peptide and the Andromeda search engine score. Residues with a score <100 and residues that were also detected in the dephosphorylated sample were excluded.

