## Supplementary material for "Interactions with multiple inner kinetochore proteins determine mitotic localization of FACT": Figs. S1-S9, Table S1

### Supplementary Materials

Figs. S1 to S9
Table S1

Fig. S1.

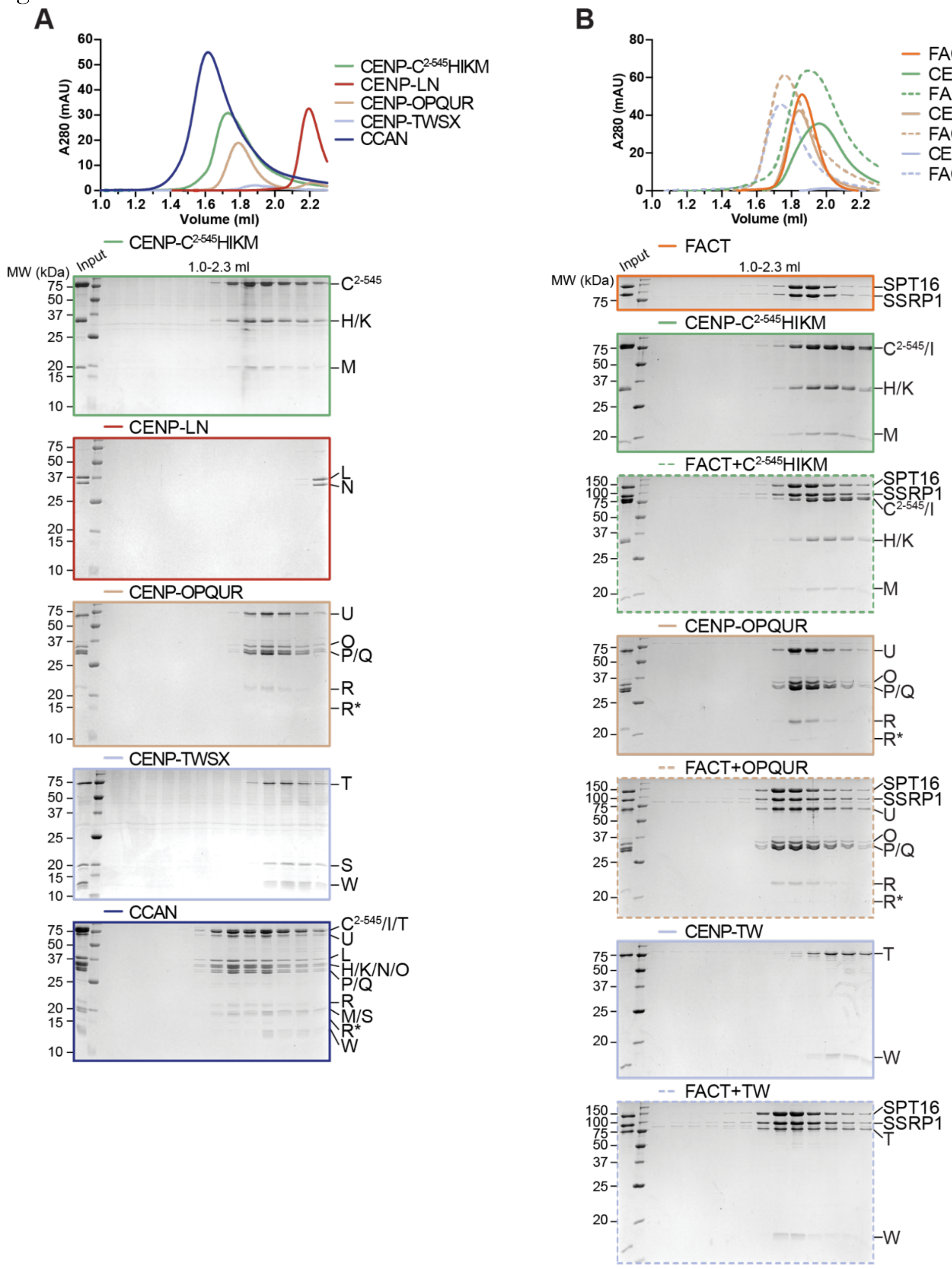

A
60
50
40
30
20
10
0
A280 (mAU)
1.0 1.2 1.4 1.6 1.8 2.0 2.2
Volume (ml)
CENP-C2-545HIKM
CENP-LN
CENP-OPQUR
CENP-TWSX
CCAN
MW (kDa)
Input
1.0-2.3 ml
C2-545
H/K
M
L
N
U
O
P/Q
R
R*
T
S
W
C2-545/I/T
H/K/N/O
M/S
B
80
FACT
CENP-C2-545HIKM
FACT+C2-545HIKM
CENP-OPQUR
FACT+OPQUR
CENP-TW
FACT+TW
SPT16
SSRP1
C2-545/I

**Figure S1 (related to Figure 1)**

**(A)** Analytical SEC of the individual CCAN subcomplexes and its reconstitution.

**(B)** Analytical SEC of FACT and CENP-$C^{2-545}$HIKM, CENP-OPQUR and CENP-TW with Coomassie-stained SDS-PAGEs below.

Fig. S2.

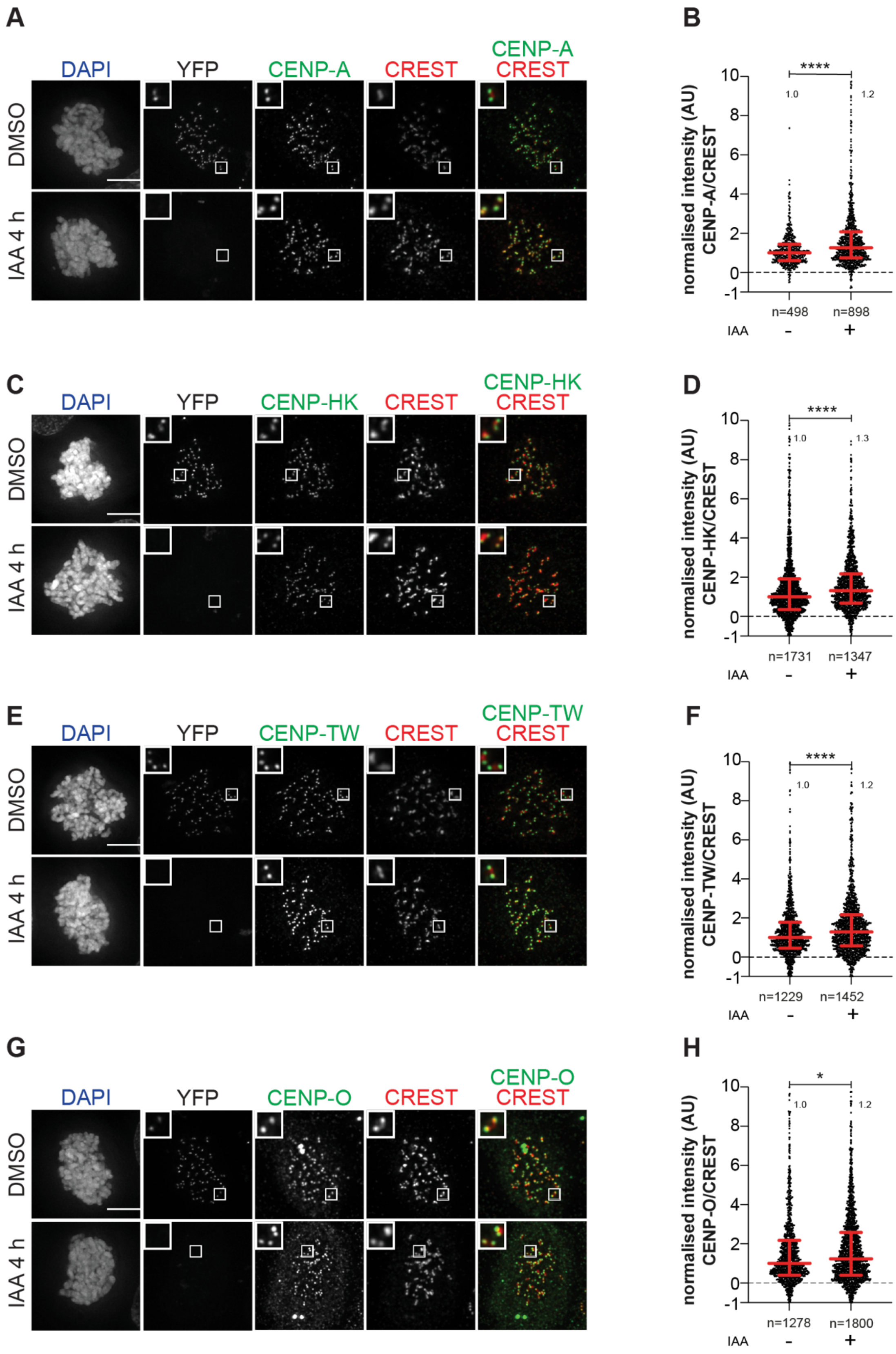
A
DAPI
YFP
CENP-A
CREST
CENP-A
CREST
DMSO
IAA 4 h
B
****
1.0
1.2
normalised intensity (AU)
CENP-A/CREST
n=498
n=898
IAA
-
+
C
DAPI
YFP
CENP-HK
CREST
CENP-HK
CREST
DMSO
IAA 4 h
D
****
1.0
1.3
normalised intensity (AU)
CENP-HK/CREST
n=1731
n=1347
IAA
-
+
E
DAPI
YFP
CENP-TW
CREST
CENP-TW
CREST
DMSO
IAA 4 h
F
****
1.0
1.2
normalised intensity (AU)
CENP-TW/CREST
n=1229
n=1452
IAA
-
+
G
DAPI
YFP
CENP-O
CREST
CENP-O
CREST
DMSO
IAA 4 h
H
*
1.0
1.2
normalised intensity (AU)
CENP-O/CREST
n=1278
n=1800
IAA
-
+

**Figure S2 (related to Figure 1)**

- **(A)** Representative images of localization of CENP-A after degradation of CENP-C in DLD-1-CENP-C$^{YFP-AID}$ cells for 4 hours. Cells were treated with IAA (500 μM) to degrade endogenous CENP-C and Nocodazole (3.3 μM) to get mitotic population of cells for 4 hours. CREST serum was used to visualize kinetochores and DAPI to stain DNA. Three biological replicates were performed. Scale bar: 5 μm.
- **(B)** Scatter plot of CENP-A levels at kinetochores for the experiment shown in panel (A). *n* is the number of individually measured kinetochores.
- **(C)** Representative images of localization of CENP-HK after degradation of CENP-C in DLD-1-CENP-C$^{YFP-AID}$ cells for 4 h. Cells were treated with IAA (500 μM) to degrade endogenous CENP-C and Nocodazole (3.3 μM) to get mitotic population of cells for 4 hours. CREST serum was used to visualize kinetochores and DAPI to stain DNA. Three biological replicates were performed. Scale bar: 5 μm.
- **(D)** Scatter plot of CENP-HK levels at kinetochores for the experiment shown in panel (C). *n* refers to individually measured kinetochores.
- **(E)** Representative images of localization of CENP-TW after degradation of CENP-C in DLD-1-CENP- C$^{YFP-AID}$ cells for 4 hours. Cells were treated with IAA (500 μM) to degrade endogenous CENP-C and with Nocodazole (3.3 μM) for 4 hours to enrich for mitotic cells. CREST serum was used to visualize kinetochores and DAPI to stain DNA. Three biological replicates were performed. Scale bar: 5 μm.
- **(F)** Scatter plot of CENP-TW levels at kinetochores for the experiment shown in panel (E). *n* refers to individually measured kinetochores.
- **(G)** Representative images of localization of CENP-O after degradation of CENP-C in DLD-1-CENP-C$^{YFP-AID}$ cells for 4 hours. Cells were treated with IAA (500 μM) to degrade endogenous CENP-C and Nocodazole (3.3 μM) to get mitotic population of cells for 4 hours. CREST was used to visualize kinetochores and DAPI to stain DNA. Three biological replicates were performed. Scale bar: 5 μm.
- **(H)** Scatter plot of CENP-O levels at kinetochores for the experiment in panel (G). *n* is the number of individually measured kinetochores.

Fig. S3.

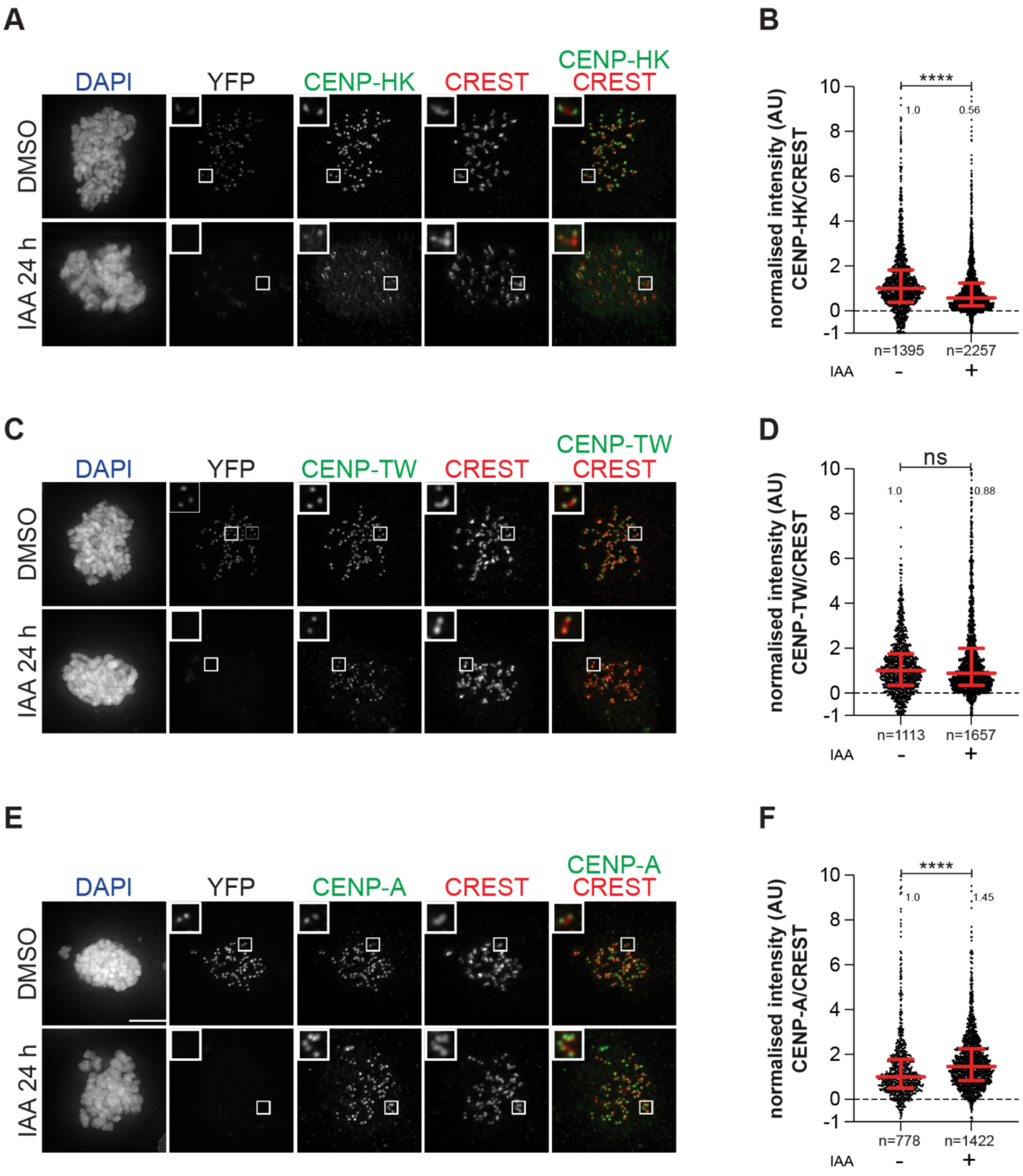


**Figure S3 (related to Figure 1)**

**(A)** Representative images of localization of CENP-HK after degradation of CENP-C in DLD-1-CENP-C$^{YFP-AID}$ cells for 24 hours. Cells were treated with IAA (500 μM) to degrade endogenous CENP-C for 24 hours and Nocodazole (3.3 μM) for 4 hours to get mitotic population of cells. CREST was used to visualize kinetochores and DAPI to stain DNA. Three biological replicates were performed. Scale bar: 5 μm.

**(B)** Scatter plot of CENP-HK levels at kinetochores of the experiment shown in panel (A). *n* is the number of individually measured kinetochores.

**(C)** Representative images of localization of CENP-TW after degradation of CENP-C in DLD-1-CENP-$C^{YFP-AID}$ cells for 24 hours. Cells were treated with IAA (500 μM) to degrade endogenous CENP-C for 24 hours and Nocodazole (3.3 μM) for 4 hours to get mitotic population of cells. CREST serum was used to visualize kinetochores and DAPI to stain DNA. Three biological replicates were performed. Scale bar: 5 μm.

**(D)** Scatter plot of CENP-TW levels at kinetochores of the experiment shown in panel (C). *n* is the number of individually measured kinetochores.

**(E)** Representative images of localization of CENP-A after degradation of CENP-C in DLD-1-CENP-$C^{YFP-AID}$ cells for 24 hours. Cells were treated with IAA (500 μM) to degrade endogenous CENP-C for 24 hours. Nocodazole (3.3 μM) was added for 4 hours to enrich for mitotic cells. CREST serum was used to visualize kinetochores and DAPI to stain DNA. Three biological replicates were performed. Scale bar: 5 μm.

**(F)** Scatter plot of CENP-A levels at kinetochores of the experiment shown in panel (E). *n* is the number of individually measured kinetochores.

**Fig. S4.**

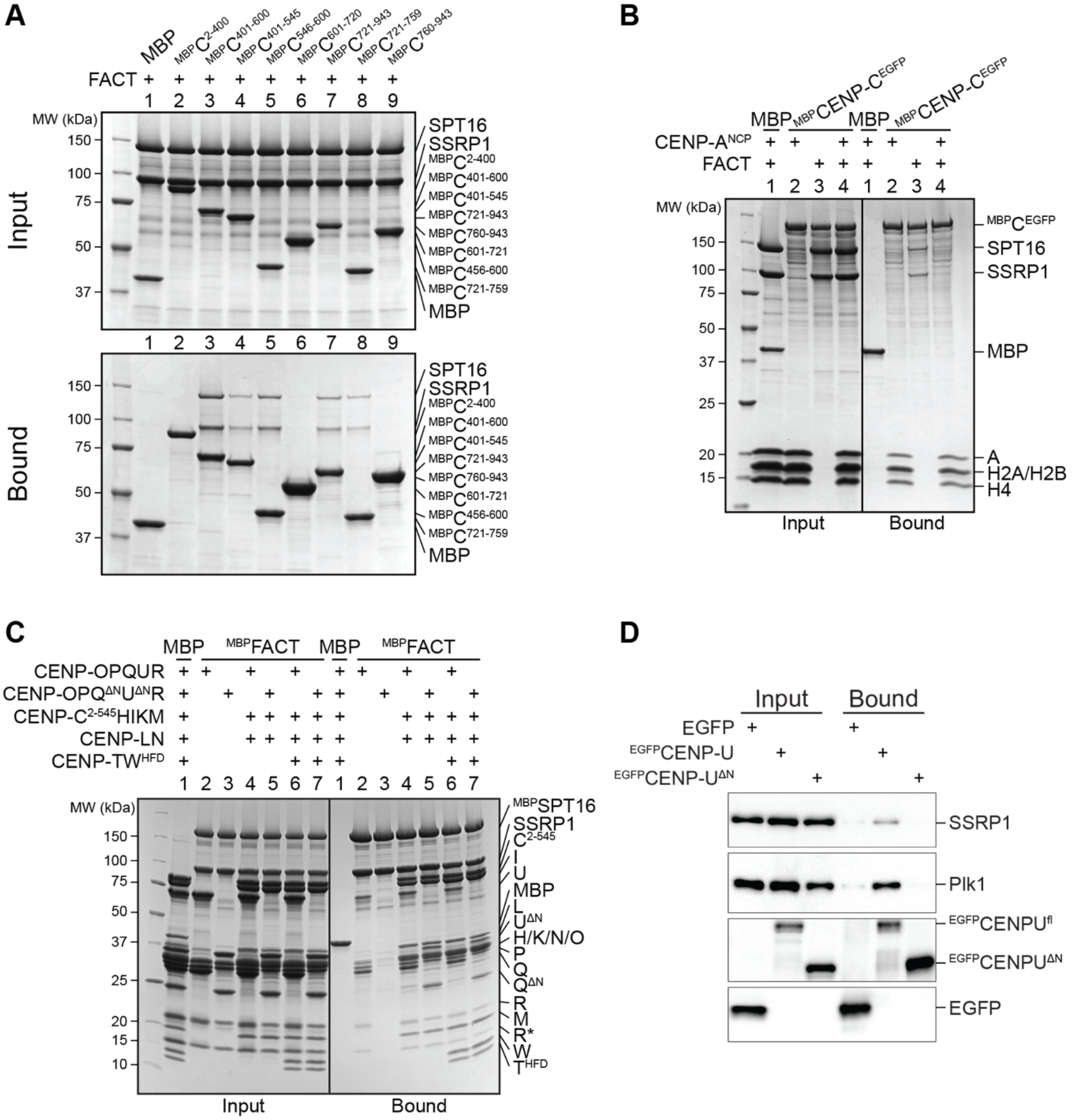


**Figure S4 (related to Figure 2)**

**(A)** Amylose-resin pull-down assay with a set of $^{MBP}$CENP-C fusion proteins as baits spanning the entire sequence of CENP-C and FACT as a prey.

**(B)** Amylose-resin pull-down assay using immobilized $^{MBP}$CENP-$C^{EGFP}$ on beads and adding FACT and CENP-$A^{NCP}$ as preys. CENP-$A^{NCP}$ is the histone octamer reconstituted on 145-bp Widom 601 sequence.

**(C)** Amylose-resin pull-down assay with $^{MBP}$FACT as a bait to analyze the influence of the CENP-QU N-terminal tails on the binding of CENP-OPQUR, -$C^{2\text{-}545}$HIKMLNOPQUR and -$C^{2\text{-}545}$HIKMLNOPQURTW$^{HFD}$.

**(D)** Lysates prepared from STLC synchronized DLD-1 cells expressing GFP alone or GFP-CENP-U Fl or 115-C were subjected to immunoprecipitation using GFP-trap beads followed by Western blotting with antibodies against GFP, SSRP1 and PLK1. PLK1 was used as internal control.

**Fig. S5.**

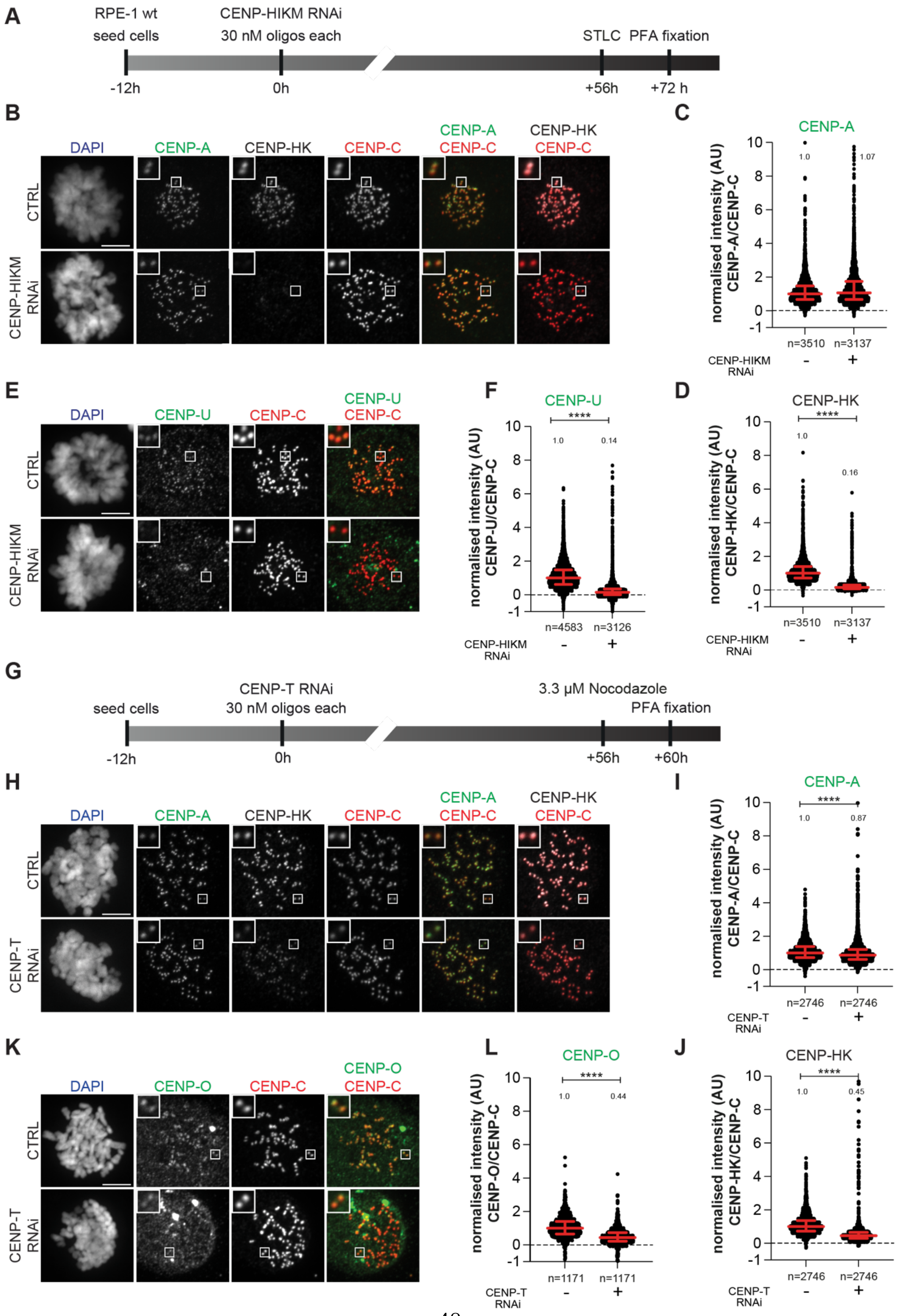

A
RPE-1 wt
seed cells
CENP-HIKM RNAi
30 nM oligos each
STLC PFA fixation
-12h
0h
+56h
+72 h
B
DAPI
CENP-A
CENP-HK
CENP-C
CENP-A CENP-C
CENP-HK CENP-C
CTRL
CENP-HIKM RNAi
C
CENP-A
normalised intensity (AU) CENP-A/CENP-C
1.0
1.07
n=3510
n=3137
CENP-HIKM RNAi
-
+
E
DAPI
CENP-U
CENP-C
CENP-U CENP-C
CTRL
CENP-HIKM RNAi
F
CENP-U
****
1.0
0.14
normalised intensity (AU) CENP-U/CENP-C
n=4583
n=3126
CENP-HIKM RNAi
-
+
D
CENP-HK
****
1.0
0.16
normalised intensity (AU) CENP-HK/CENP-C
n=3510
n=3137
CENP-HIKM RNAi
-
+
G
seed cells
CENP-T RNAi
30 nM oligos each
3.3 μM Nocodazole
PFA fixation
-12h
0h
+56h
+60h
H
DAPI
CENP-A
CENP-HK
CENP-C
CENP-A CENP-C
CENP-HK CENP-C
CTRL
CENP-T RNAi
I
CENP-A
****
1.0
0.87
normalised intensity (AU) CENP-A/CENP-C
n=2746
n=2746
CENP-T RNAi
-
+
K
DAPI
CENP-O
CENP-C
CENP-O CENP-C
CTRL
CENP-T RNAi
L
CENP-O
****
1.0
0.44
normalised intensity (AU) CENP-O/CENP-C
n=1171
n=1171
CENP-T RNAi
-
+
J
CENP-HK
****
1.0
0.45
normalised intensity (AU) CENP-HK/CENP-C
n=2746
n=2746
CENP-T RNAi
-
+

**Figure S5 (related to Figure 3)**

**(A)** Schematic representation of experimental scheme used for CENP-HIKM RNAi

**(B)** Representative images of localization of CENP-A and CENP-HK after depletion of CENP-HIKM complex in RPE-1 cells. CENP-HIKM RNAi was performed for 72 hours using oligos for each subunit at 30 nM concentration. Cells were treated with STLC (5 μM) for 16 hours to obtain a mitotic population before fixation. CENP-C was used to visualize kinetochores and DAPI to stain DNA. Three biological replicates were performed. Scale bar: 5 μm.

**(C)** Scatter plot of CENP-A levels at kinetochores for the experiment shown in panel (B). *n* is the number of individually measured kinetochores.

**(D)** Scatter plot of CENP-HK levels at kinetochores for the experiment in panel (B). *n* is the number of individually measured kinetochores.

**(E)** Representative images of localization of CENP-U after depletion of CENP-HIKM complex in RPE-1 cells. CENP-HIKM RNAi was performed for 72 hours using oligos for each subunit at 30 nM concentration. To obtain mitotic cells, cells were treated with STLC (5 μM) for 16 hours before fixation. CENP-C visualizes kinetochores and DAPI stains DNA. Three biological replicates were performed. Scale bar: 5 μm.

**(F)** Scatter plot of CENP-U levels at kinetochores of the experiment shown in panel (E). *n* refers to individually measured kinetochores.

**(G)** Schematic representation of experimental scheme used for CENP-T RNAi

**(H)** Representative images of localization of CENP-A and CENP-HK after depletion of CENP-T complex in RPE-1 cells. CENP-T RNAi was performed for 60 hours using oligos for each subunit at 30 nM concentration. Cells were treated with Nocodazole (3.3 μM) for 4 hours before fixation to enrich for mitotic cells. CENP-C visualizes kinetochores and DAPI stains DNA. Three biological replicates were performed. Scale bar: 5 μm.

**(I)** Scatter plot of CENP-A levels at kinetochores for the experiment in panel (H). *n* is the number of individually measured kinetochores.

**(J)** Scatter plot of CENP-HK levels at kinetochores for the experiment in panel (H). *n* is the number of individually measured kinetochores.

**(K)** Representative images of localization of CENP-O after depletion of CENP-T complex in RPE-1 cells. CENP-T RNAi was performed for 60 hours using oligos for each subunit at 30 nM concentration. Cells were treated with Nocodazole (3.3 μM) for 4 hours before fixation to obtain mitotic population. CENP-C visualizes kinetochores and DAPI stains DNA. Three biological replicates were performed. Scale bar: 5 μm.

**(L)** Scatter plot of CENP-O levels at kinetochores for the experiment in panel (K). *n* is the number of individually measured kinetochores.

**Fig. S6.**

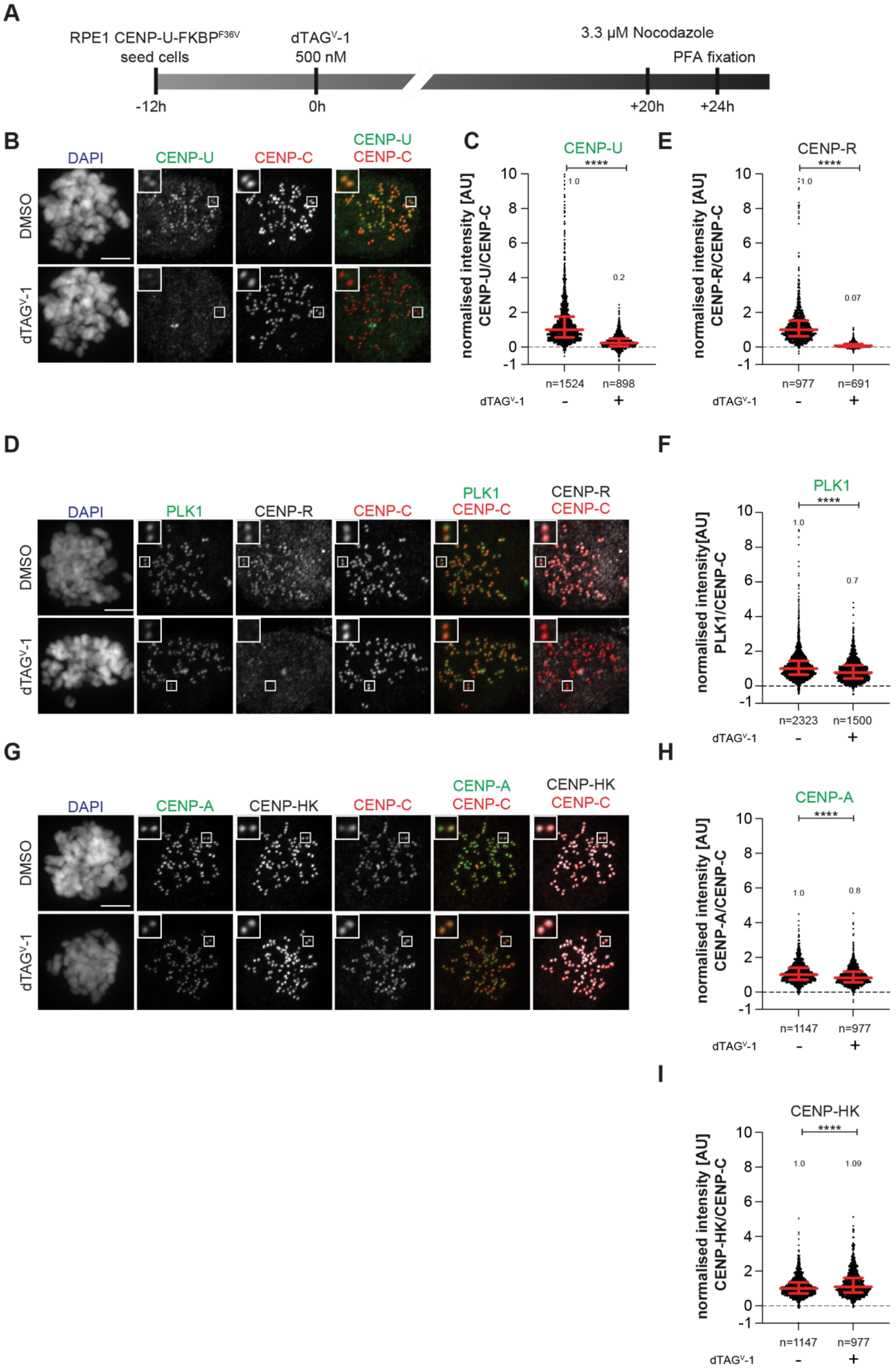
A
RPE1 CENP-U-FKBPF36V
seed cells
dTAGV-1
500 nM
3.3 μM Nocodazole
PFA fixation
-12h
0h
+20h
+24h
B
DAPI
CENP-U
CENP-C
CENP-U
CENP-C
DMSO
dTAGV-1
C
CENP-U
normalised intensity [AU]
CENP-U/CENP-C
****
1.0
0.2
n=1524
n=898
dTAGV-1
-
+
E
CENP-R
normalised intensity [AU]
CENP-R/CENP-C
****
1.0
0.07
n=977
n=691
D
DAPI
PLK1
CENP-R
CENP-C
PLK1
CENP-C
CENP-R
CENP-C
F
PLK1
normalised intensity[AU]
PLK1/CENP-C
****
1.0
0.7
n=2323
n=1500
G
DAPI
CENP-A
CENP-HK
CENP-C
CENP-A
CENP-C
CENP-HK
CENP-C
H
CENP-A
normalised intensity [AU]
CENP-A/CENP-C
****
1.0
0.8
n=1147
n=977
I
CENP-HK
normalised intensity [AU]
CENP-HK/CENP-C
****
1.0
1.09
n=1147
n=977

**Figure S6 (related to Figure 3)**

**(A)** Schematic representation of experimental scheme used for CENP-U dTAG$^{V}$-1 treatment

**(B)** Representative images of localization of CENP-U after depletion of CENP-U complex in RPE-1-CENP-U-FKBP$^{F36V}$ cells. Cells were treated with dTAG$^{V}$-1 (500 nM) for 24 hours to degrade endogenous CENP-U complex. Cells were treated with Nocodazole (3.3 μM) for 4 hours to obtain a mitotic cell population prior to fixation. CENP-C was used to visualize kinetochores and DAPI to stain DNA. Three biological replicates were performed. Scale bar: 5 μm.

**(C)** Scatter plot of CENP-U levels at kinetochores for the experiment in panel (B). *n* is the number of individually measured kinetochores.

**(D)** Representative images of localization of CENP-R and PLK1 after depletion of CENP-U complex in RPE-1-CENP-U-FKBP$^{F36V}$ cells. Cells were treated with dTAG$^{V}$-1 (500 nM) for 24 hours to degrade endogenous CENP-U complex. Cells were treated with Nocodazole (3.3 μM) for 4 h prior to fixation to obtain mitotic population. CENP-C was used to visualize kinetochores and DAPI to stain DNA. Three biological replicates were performed. Scale bar: 5 μm.

**(E)** Scatter plot of CENP-R levels at kinetochores for the experiment in panel (D). *n* is the number of individually measured kinetochores.

**(F)** Scatter plot of PLK1 levels at kinetochores for the experiment in panel (D). *n* is the number of individually measured kinetochores.

**(G)** Representative images of localization of CENP-A and CENP-HK after depletion of CENP-U complex in RPE-1-CENP-U-FKBP$^{F36V}$ cells. Cells were treated with dTAG$^{V}$-1 (500 nM) for 24 hours to degrade endogenous CENP-U complex. Cells were treated with Nocodazole (3.3 μM) for 4 h prior to fixation to obtain mitotic population. CENP-C was used to visualize kinetochores and DAPI to stain DNA. Three biological replicates were performed. Scale bar: 5 μm.

**(H)** Scatter plot of CENP-A levels at kinetochores of the experiment shown in panel (G). *n* is the number of individually measured kinetochores.

**(I)** Scatter plot of CENP-HK levels at kinetochores of the experiment shown in panel (G). *n* is the number of individually measured kinetochores.

Fig. S7.

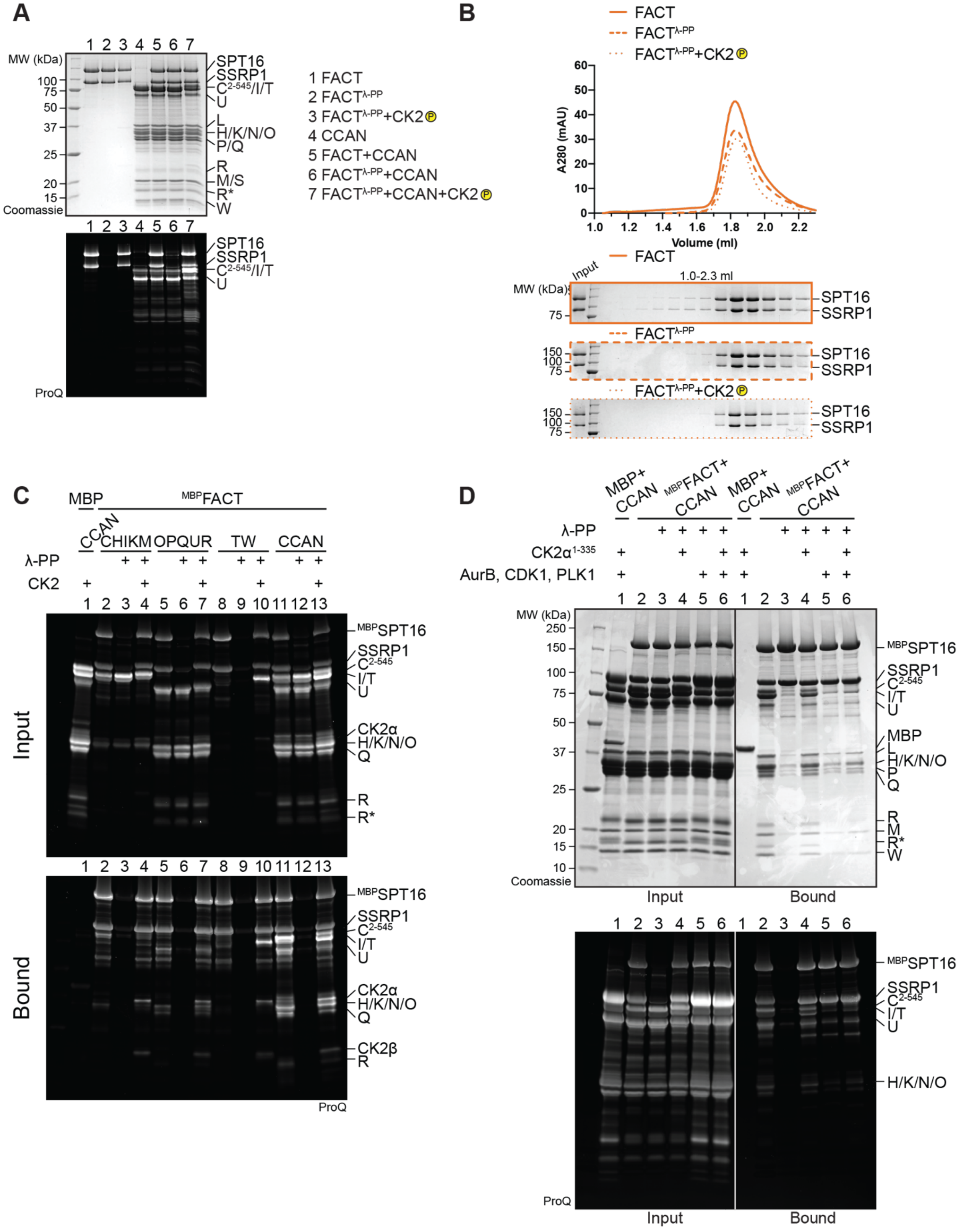
A
MW (kDa)
1 2 3 4 5 6 7
100
75
50
37
25
20
15
Coomassie
SPT16
SSRP1
C2-545/I/T
U
L
H/K/N/O
P/Q
R
M/S
R*
W
ProQ
1 FACT
2 FACTλ-PP
3 FACTλ-PP+CK2
4 CCAN
5 FACT+CCAN
6 FACTλ-PP+CCAN
7 FACTλ-PP+CCAN+CK2
B
FACT
FACTλ-PP
FACTλ-PP+CK2
A280 (mAU)
Volume (ml)
Input
1.0-2.3 ml
MW (kDa)
150
100
75
SPT16
SSRP1
C
MBP
MBPFACT
CCAN
CHIKM
OPQUR
TW
CCAN
λ-PP
CK2
1 2 3 4 5 6 7 8 9 10 11 12 13
Input
Bound
MBPSPT16
SSRP1
C2-545
I/T
U
CK2α
H/K/N/O
Q
R
R*
CK2β
ProQ
D
MBP+ CCAN
MBPFACT+ CCAN
λ-PP
CK2α1-335
AurB, CDK1, PLK1
MW (kDa)
250
150
100
75
50
37
25
20
15
10
Coomassie
Input
Bound
MBPSPT16
SSRP1
C2-545
I/T
U
MBP
L
H/K/N/O
P
Q
R
M
R*
W
ProQ

**Figure S7 (related to Figure 5)**

**(A)** ProQ Diamond staining to monitor the phosphorylation state of samples in Figure 5A.

**(B)** Additional samples of the analytical SEC experiment in Figure 5A verify that the phosphorylation state of FACT does not alter its elution volume.

**(C)** ProQ Diamond staining of the pull-down assay in Figure 5B.

**(D)** Amylose-resin pull-down assay to analyze CCAN binding to dephosphorylated $^{MBP}$FACT upon phosphorylation by CK2α$^{1\text{-}335}$ or Aurora B, CDK1 and PLK1 or all of them. CDK1 indicates the use of a complex of CDK1/Cyclin-B/CKS1. The ProQ Diamond staining of the SDS-PAGE is shown below.

Fig. S8.

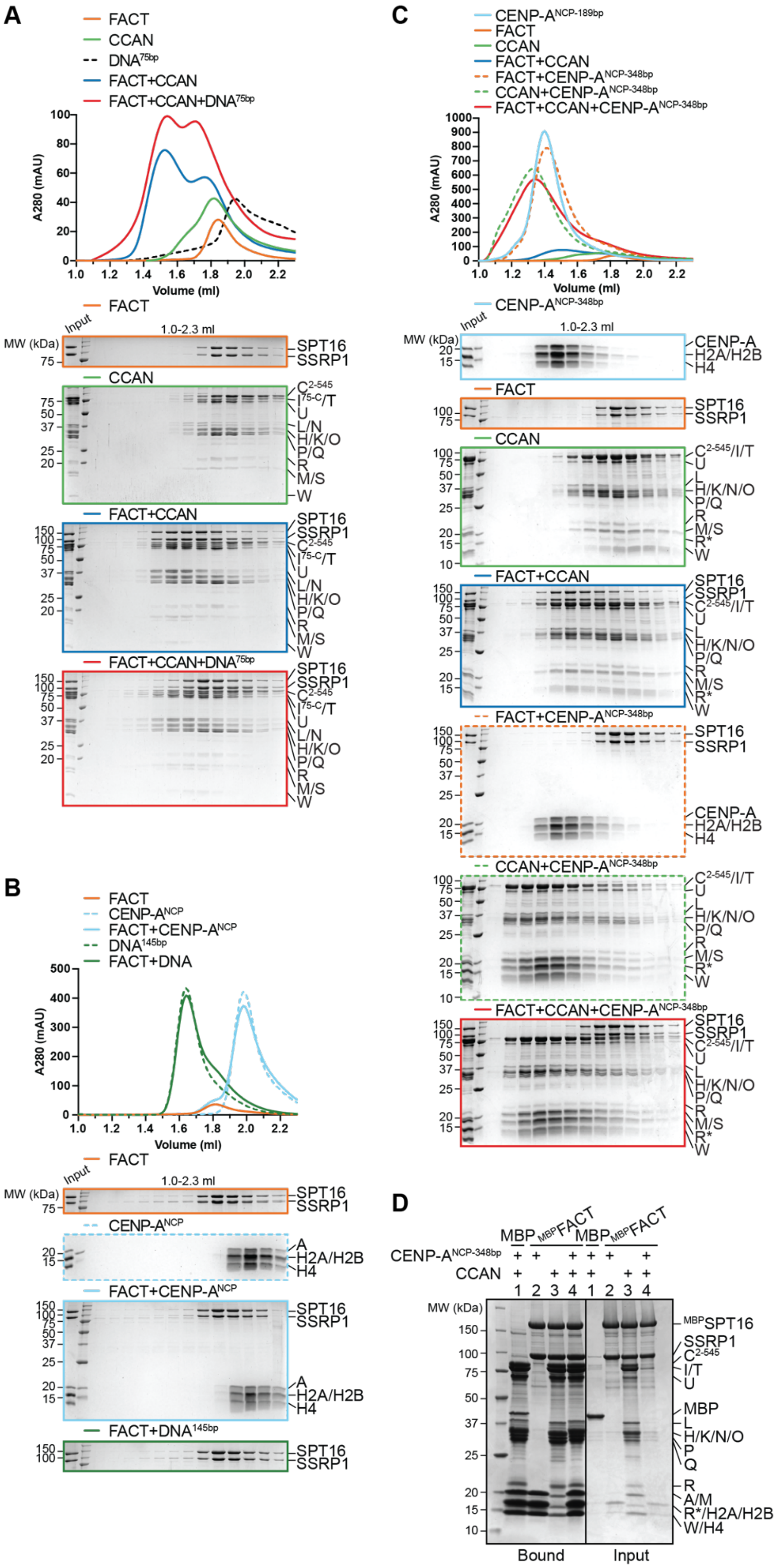

A
FACT
CCAN
DNA75bp
FACT+CCAN
FACT+CCAN+DNA75bp
A280 (mAU)
Volume (ml)
MW (kDa)
Input
1.0-2.3 ml
SPT16
SSRP1
C2-545
I75-C/T
U
L/N
H/K/O
P/Q
R
M/S
W
B
FACT
CENP-ANCP
FACT+CENP-ANCP
DNA145bp
FACT+DNA
FACT+DNA145bp
A
H2A/H2B
H4
C
CENP-ANCP-189bp
FACT+CENP-ANCP-348bp
CCAN+CENP-ANCP-348bp
FACT+CCAN+CENP-ANCP-348bp
CENP-ANCP-348bp
CENP-A
C2-545/I/T
H/K/N/O
R*
D
MBP
MBPFACT
CCAN
MBPSPT16
I/T
MBP
A/M
R*/H2A/H2B
W/H4
Bound
Input

**Figure S8 (related to Figure 6)**

**(A)** Analytical SEC of FACT and CCAN upon the addition of a 75-bp CEN1-like DNA.

**(B)** Analytical SEC to test binding of FACT to CENP-$A^{NCP}$ or 145-bp Widom 601 DNA.

**(C)** Analytical SEC of FACT, CCAN, and CENP-$A^{NCP}$ on a 348-bp DNA. The histone octamer was reconstituted on a 183-bp CEN1-like sequence and 165 CEN1-like sequence was ligated to it.

**(D)** Amylose-resin pull-down assay of $^{MBP}$FACT and CCAN upon addition of CENP-$A^{NCP\text{-}348bp}$.

**Fig. S9.**

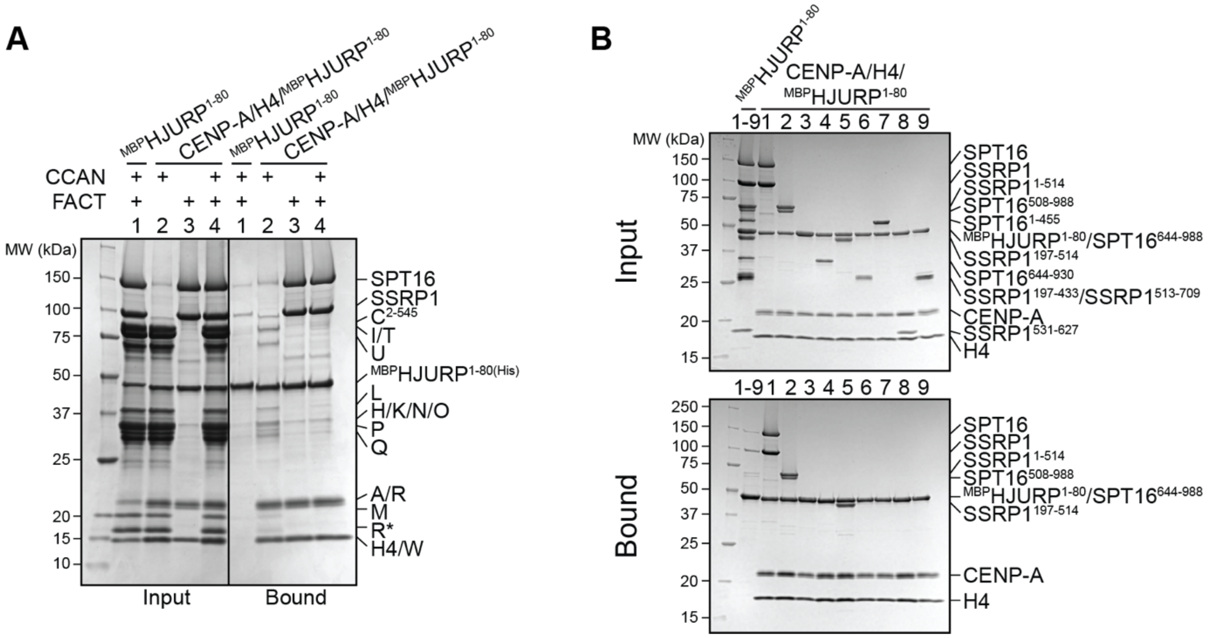


**Figure S9 (related to Figure 6)**

**(A)** Amylose-resin pull-down assay using $^{MBP}$HJURP$^{1-80}$ in complex with CENP-A/H4 as bait and CCAN and FACT as preys. $^{MBP}$HJURP$^{1-80}$ in absence of histones was used as a negative control.

**(B)** Amylose-resin pull-down assay using $^{MBP}$HJURP$^{1-80}$ in complex with CENP-A/H4 as bait and FACT constructs (Fig. 4A) as preys. $^{MBP}$HJURP$^{1-80}$ in absence of histones was used as a negative control.

**Table S1.**

| | Position | Phospho (STY) Probabilities | PEP | Score |
|---|---|---|---|---|
| **SPT16** | 979; 986 | E**pS**(0.977)LGS(0.023)EEE **pS**(1)GKDWDELEEEAR | 4.29E-22 | 170.53 |
| | 1004 | ADRE**pS**(1)RYEEEEEQSR | 2.96E-13 | 190.55 |
| **SSRP1** | 437 | EGMNP**pS**(0.731)Y(0.093)DEYAD **pS**(1)DEDQHDAYLER | 2.97E-16 | 160.43 |
| | 438 | EGMNPS(0.292)**pY**(0.707)DEY(0.002) AD**pS**(0.999)DEDQHDAYLER | 7.50E-09 | 112.99 |
| | 444 | EGMNPSYDEYAD**pS**(1)DEDQHDAYLER | 2.65E-106 | 389.53 |
| | 667 | EFV**pS**(0.999)S(0.001)DESSSGENK | 5.17E-29 | 265.89 |

Phospho-sites on FACT detected by mass spectrometry subsequent to treatment with CK2. Sites in bold followed by the probability, the p-value of the posterior error probability (PEP) of the peptide and the Andromeda search engine score. Residues with a score <100 and residues that were also detected in the dephosphorylated sample were excluded.
